# Omics driven onboarding of the carotenoid producing red yeast *Xanthophyllomyces dendrorhous* CBS 6938

**DOI:** 10.1101/2023.07.31.551333

**Authors:** Emma E. Tobin, Joseph H. Collins, Celeste B. Marsan, Gillian T. Nadeau, Kim Mori, Anna Lipzen, Stephen Mondo, Igor V. Grigoriev, Eric M. Young

**Affiliations:** Life Sciences and Bioengineering Center, Department of Chemical Engineering, Worcester Polytechnic Institute, Worcester, MA; U.S. Department of Energy Joint Genome Institute, Lawrence Berkeley National Laboratory, Berkeley, CA 94720; Plant and Microbial Biology, University of California Berkeley, Berkeley, CA 94720

**Author notes:** Equal contribution (Emma E. Tobin, Joseph H. Collins).

**Keywords:** *X.dendrorhous*, transcriptomics, regulation, DGE, nonconventional, photobiology

## Abstract

Transcriptomics is a powerful approach for functional genomics and systems biology, yet it can also be used for genetic part discovery. Genetic part discovery has never been more necessary, as advances in synthetic biology increase the number of tractable organisms that need tunable gene expression for genetic circuits and metabolic pathways. Therefore, approaches are needed to assess a tractable organism and obtain a convenient set of genetic parts to support future research. Here, we describe a genomic and transcriptomic approach to derive a modular integrative part library with constitutive and regulated promoters in the basidiomycete yeast Xanthophyllomyces dendrorhous CBS 6938. X. dendrorhous is currently the sole biotechnologically relevant organism in the Tremellomycete family - it produces large amounts of astaxanthin, especially under oxidative stress and exposure to light. Particularly for this yeast, there are not large libraries of parts from related organisms that could be transferred. They must be derived. To do this, a contiguous genome was first obtained through combined short read and long read sequencing. Then, differential gene expression (DGE) analysis using transcriptomics was performed, comparing oxidative stress and exposure to different wavelengths of light. This revealed a set of putative light-responsive regulators that mediate a complex survival response to ultraviolet (UV) where X. dendrorhous upregulates aromatic amino acid and tetraterpenoid biosynthesis and downregulates central carbon metabolism and respiration. The DGE data was then used to derive 26 constitutive and regulated gene expression elements from the genome. The gene expression elements were designed to be compatible with a new modular cloning system for X. dendrorhous which includes integration sites, terminators, selection markers, and reporters. Each element was characterized by luciferase assay of an integrated gene expression cassette. Notably, a novel promoter from a hypothetical gene that has 9-fold activation upon UV exposure was characterized. This study defines an advanced modular genetic part collection for engineering the basidiomycete X. dendrorhous CBS 6938 while simultaneously discovering potential targets for increasing tetraterpenoid biosynthesis. Further, it demonstrates that -omics-to-parts workflows can simultaneously provide useful genomic data and advance genetic tools for nonconventional microbes, particularly those without a related model organism. This approach will be broadly useful in current efforts to engineer diverse microbes.

**KEY POINTS:**

- Omics-to-parts can be applied to non-model organisms for rapid “onboarding”.
- 26 promoters native to *X. dendrorhous* were identified.
- Omics revealed unique photobiology in *X. dendrorhous*.

## INTRODUCTION

Yeasts are a diverse group of fungi characterized by unicellular growth, which is useful in industrial fermentations. The group is not monophyletic, rather the yeast state has emerged several times across the fungal kingdom (Nagy et al 2014). Therefore, yeasts have different genome architectures, metabolic capabilities, and stress tolerances that are not only due to the ecological niches to which they are adapted, but also due to their different lineages (Boekhout et al 2021). The impact this biodiversity might have on synthetic biology and industrial biotechnology is currently being explored (Wagner & Alper 2016; Steensels et al. 2014; Steensels & Verstrepen 2014; Snoek et al. 2015). However, the different lineages limit the transferability of genetic tools between yeasts (Zeevi et al. 2014; Cao et al. 2017), particularly for the basidiomycete yeasts that are not closely related to any model organism (Wen et al. 2020; Otoupal et al. 2019; Johnson 2013). Genomics analysis of the fungal kingdom is well-developed (Almeida et al. 2014; Libkind et al. 2020; Peris et al. 2023; Collins et al. 2021; Merényi et al. 2023), which is a foundation that can be leveraged to derive a collection of genetic parts in addition to enabling insight into fundamental biology. Here, we demonstrate an -omics-to-parts workflow for the basidiomycete yeast *Xanthophyllomyces dendrorhous* [*Phaffia rhodozyma*], currently the only biotechnologically relevant organism from the Tremellomycete family of Basidiomycota (Bellora et al. 2016).

The model yeast, the ascomycete *Saccharomyces cerevisiae*, has long been the host of choice for yeast metabolic engineering (Da Silva et al. 2012; Jeppsson et al. 2003; Shen et al. 2012; Farhi et al. 2011; Young et al. 2018; Liu et al. 2024; Galanie et al. 2015; Hong & Nielsen et al. 2012). While valuable for converting hexoses to ethanol, engineering *S. cerevisiae* to consume other carbon sources and produce other economically valuable compounds can require extensive reprogramming, which imposes metabolic and gene expression burden (Li et al. 2022; Lie et al. 2013; Karim et al. 2013). This is particularly true of tolerance, which can be due to many genes interacting in currently unpredictable ways. This has led to a growing body of literature that investigates other yeasts, termed nonconventional simply because they are not *S. cerevisiae*, as potential hosts for metabolic engineering since they are already adapted to consume desired carbon sources, produce desired classes of molecules, and tolerate different fermentation conditions, thus less burden is incurred when engineering them (Torres-Haro et al. 2021; Rodríguez-Sáiz et al. 2010; Yaguchi et al. 2017; Pyne et al. 2023; Löbs et al. 2017). Methods for developing genetic tools are needed for these organisms.

Developing genomics and genetic tools in nonconventional yeasts is challenging. This is because a high-quality genome, gene regulatory data, species-specific DNA parts for genetic manipulation, and efficient transformation procedures are often missing. The process of developing these elements, termed “onboarding,” can be accelerated for nonconventional ascomycete yeasts closely related to *S. cerevisiae* because the genomics and genetics are similar, permitting transfer of similar promoter and plasmid designs (Durmusoglu et al. 2023). Onboarding organisms from relatively unknown classes, particularly basidiomycete yeasts, does not have this advantage because there is not a closely related model organism with a complete, contiguous genome and well-developed genetics.

*Xanthophyllomyces dendrorhous* is a nonconventional yeast that has gained attention in the aquaculture (Kheirabadi et al. 2022; Bjerkeng et al. 2007; Bampidis et al. 2022) and nutraceutical (Ambati et al. 2014; Wang et al. 2019; Gimeno-Pérez et al. 2015; Fernández-Arrojo et al. 2007) industries due to its natural ability to produce astaxanthin, a vibrant red pigment with potent antioxidant properties (Naguib 2000; Higuera-Ciapara et al. 2006; Zhang et al. 2020). It is currently the only Tremellomycete with biotechnology applications. Because of its industrial potential, *X. dendrorhous* research has been centered around response to stressors (Santopietro et al. 1998; Schroeder et al. 1995; Rodríguez-Sáiz et al. 2010), ability to utilize low-cost carbon sources (Villegas-Méndez et al. 2021; Phoproek et al. 2021; Lai et al. 2023; Gervasi et al. 2018; Dominguez-Bocanegra & Torres-Munoz 2004; Ananda & Vadlani 2011; An et al. 2001), mutagenesis (Ukibe et al. 2008; Palágyi et al. 2006; Gassel et al. 2013; Chumpolkulwong et al. 1997; Baeza et al. 2009; Ang et al. 2019; An et al. 1989), astaxanthin extraction techniques (Yin et al. 2013; Mussagy et al. 2022; Batghare et al. 2018; Aguilar-Machado et al. 2020), and characterization of carotenogenic-adjacent pathways and enzymes (Ojima et al. 2006; Leiva et al. 2015; Gutiérrez et al. 2015; Córdova et al. 2017; An et al. 1999; Alcaíno et al. 2008). However, genetic engineering is limited, with the most common method being overexpression of carotenogenic genes through the increase of gene copy number (Verdoes et al. 2003; Ledetzky et al. 2014; Breitenbach et al. 2011; Gassel et al. 2014).

An organism to be used as a host for synthetic biology and precision genome engineering needs a high-quality genome sequence, a modular gene expression parts collection, and efficient transformation methods. While *X. dendrorhous* currently has several genomes sequenced (Sharma et al. 2015; Bellora et al. 2016) and successful transformation protocols (Yamamoto et al. 2016; Wery et al. 1997), there is not a high-quality contiguous genome for precision genome engineering nor is there a modular gene expression part collection. So far, only 8 constitutive promoters from central carbon metabolism are available (Rodríguez-Sáiz et al. 2009; Hara et al. 2014). Furthermore, complex but uncharacterized regulation of the astaxanthin biosynthetic pathway limits utilization of this organism as a platform for production (Rodríguez-Sáiz et al. 2010; Zhang et al. 2019; Tropea et al. 2013; Stachowiak 2013; Schroeder & Johnson 1995; Li et al. 2023; Liu et al. 2006; Huang et al. 2022; Kim et al. 2005). Therefore, a combined genomics and transcriptomics effort could reveal metabolic regulation and yield additional promoters for future engineering, which could be key to realizing the potential of this yeast.

Here, we describe an -omics-driven workflow from genome sequencing and transcriptomics to a modular gene expression parts collection for *X. dendrorhous*. The final parts collection contains 26 promoters within a hierarchical TypeIIS enzyme assembly strategy that includes terminators, selection markers, reporters, and integration cassettes. Analysis of the transcriptomics data also generates new insights into the *X. dendrorhous* response to light and oxidative stress. Thus, this work expands the understanding of *X. dendrorhous* photobiology and genetic tools available for this yeast. Therefore, this combined systems and synthetic biology approach can deliver a suite of data and tools for nonconventional yeast that have no closely related model host.

## RESULTS

### High quality contiguous genome of *X. dendrorhous* CBS 6938

A high-quality *X. dendrorhous* CBS 6938 genome was obtained through Prymetime sequencing (Collins et al. 2021). Prymetime assembled the genome into sixteen contigs over 1,000 bp long, and relative lengths of the contigs are depicted in Figure S1a. Subsequent benchmarking universal single-copy ortholog (BUSCO) analysis revealed that the CBS 6938 genome contains 94.0% complete and single copy BUSCOs (Figure S1b). This is improved over previously available genomes in contiguity and completeness.

### Foundational genetic tools for onboarding *X. dendrorhous*

In addition to obtaining an accurate genome sequence, we performed a set of experiments to make *X. dendrorhous* easier to engineer. We first examined possible selection markers. Rather than seek to create synthetic auxotrophy, which has a variety of metabolic impacts (Mülleder et al. 2012), we screened for antibiotic sensitivity beyond the two antibiotics commonly used in *X. dendrorhous -* geneticin and hygromycin (Gassel et al. 2013; Yamamoto et al. 2016; Wery et al. 1997). Specifically, we compared *X. dendrorhous* sensitivity to nourseothricin and zeocin with the common antibiotics in a minimum inhibitory concentration (MIC) assay (Methods). The results of the MIC assay are shown in Figure 1a. The most effective compound was found to be nourseothricin with 11.6% cell survival at a concentration of 10 µg/mL. Thus, we synthesized a *X. dendrorhous* codon optimized version of the *natR* gene for selection in subsequent engineering, termed *XdNatR*.

**Figure 1.**
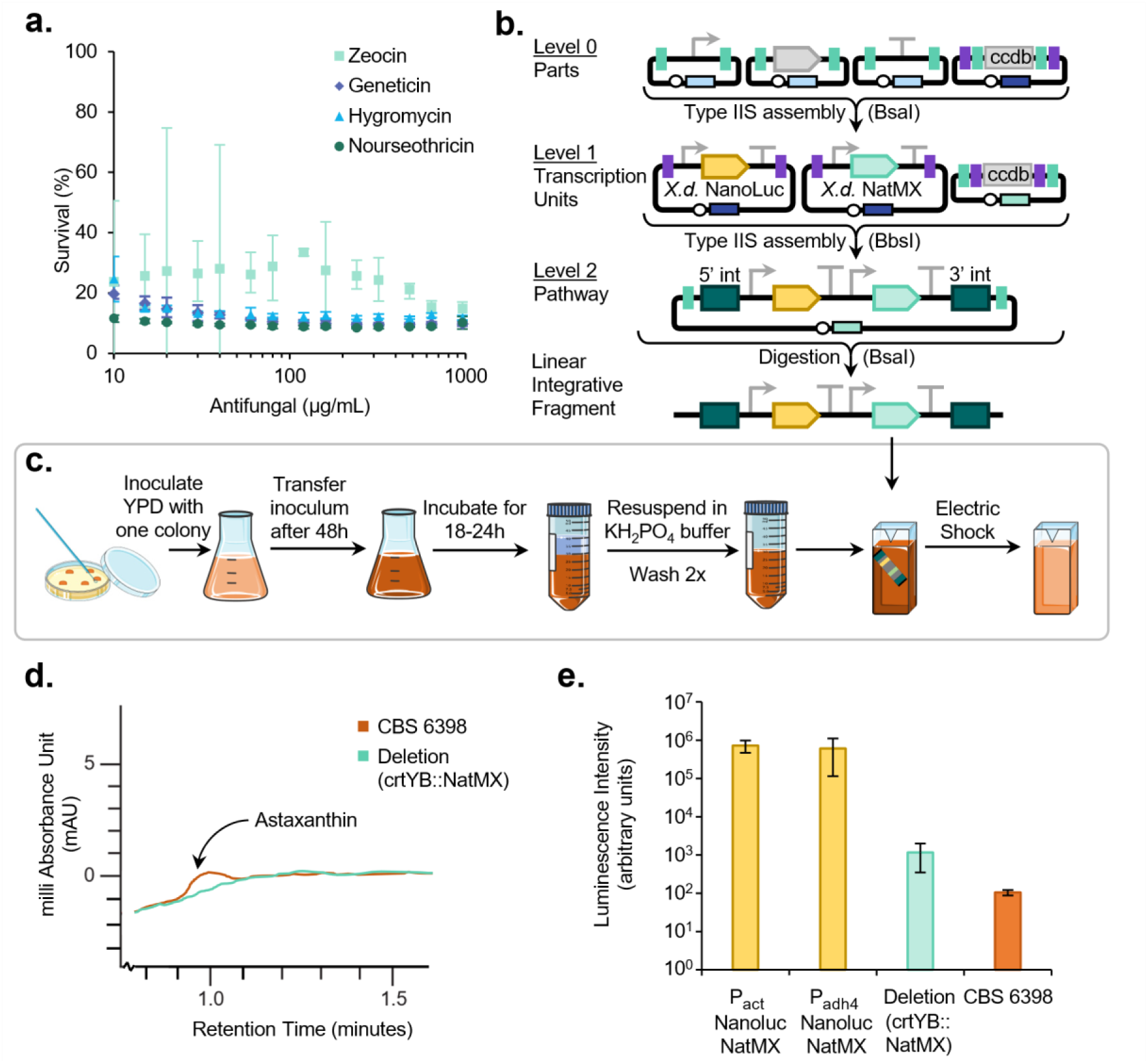
Improving foundational *X. dendrorhous* genetic tools. **(a)** Minimum Inhibitory Concentration (MIC) results depicting the survival percentage of *X. dendrorhous* CBS 6398 when exposed to the antibiotics zeocin, geneticin, hygromycin, and nourseothricin. **(b)** Hierarchical TypeIIS enzyme assembly strategy to generate fragments for homologous recombination into the *crtYB* site. **(c)** BioRender depiction of the high-efficiency transformation procedure. **(d)** Confirmation of deletion of *crtYB* activity via UHPLC. Wild type is shown in red and the *crtYB* knockout is shown in black. Overlapped UHPLC chromatograms show the lack of astaxanthin at a retention time of 0.998 in the strain with the XdNatMX cassette inserted. **(e)** Luminescence of two strains expressing Nanoluc with strong constitutive promoters, P_act_ and P_adh4_, the *X. dendrorhous* crtYB::NatMX deletion, and the *X. dendrorhous* CBS 6938 wild type. *Abbreviations:* Nanoluc - nanoluciferase, NatMX - nourseothricin resistance transcription unit.

We then designed a modular cloning system for rapidly assembling *X. dendrorhous* genetic elements (Figure 1b). This system uses the same enzymes and scars as previously published modular cloning systems based on TypeIIS restriction enzymes, often termed GoldenGate assembly (Young et al. 2018; Iverson et al. 2016). Like those systems, the *X. dendrorhous* system consists of Level 0 genetic elements (individual promoters, genes, and terminators), Level 1 transcription units (combinations of promoters, genes, and terminators), and Level 2 integrative plasmids that have homology arms and the ability to integrate several transcription units along with a selection marker. Yet, there are two significant differences. The first is that the *lacZ* element in the destination plasmids is replaced with the lethal gene *ccdb* to shift from blue/white screening of correct clones to a system where only correct clones grow on a selection plate (Bernard et al. 1994). The second is that the homology arms for *S. cerevisiae* were replaced with *X. dendrorhous* homology arms flanking the *crtYB* gene derived from the resequenced genome (Table S1). Since the *crtYB* gene product catalyzes the first committed step in visibly colorful terpenoid production, replacement of *crtYB* creates a facile red/white screening method for correctly integrated clones.

To optimize the transformation protocol, we designed an initial integrative element using the TypeIIS assembly scheme. We combined the known glutamate dehydrogenase promoter (P_gdh_) and the glycerol-3-phosphate dehydrogenase terminator (T_gpd_) to express *XdNatR*, creating an XdNatMX resistance cassette, which was subsequently cloned into the integrative plasmid and transformed into *X. dendrorhous*. In our hands, the efficiency of the published transformation was low (Wery et al. 1998), so several modifications were made as described in Methods and shown in Figure 1c. Ultra-high-performance liquid-chromatography (UPLC) was then used to demonstrate successful *crtYB* knock-out through measurement of astaxanthin. Astaxanthin peaks were detected in wild type but not knock-out extractions (Figure 1d). Thus, with the design of the integration cassette and confirmation of the transformation protocol, we were able to achieve efficient and targeted integration of DNA into the *X. dendrorhous* genome using homologous recombination.

We then cloned a luminescent reporter, NanoLuc (England et al. 2016), under the control of two of the known strongest promoters, P_act_ and P_adh4_, into our modular cloning scheme along with XdNatMX (Figure 1b), integrated it into the genome, and measured expression via luciferase assay (Figure 1e). NanoLuc-producing cells created approximately 1,000-fold greater luminescence compared to the *crtYB* knockout strain, and nearly 10,000-fold greater luminescence compared to the parent strain. The higher luminescence observed from the *crtYB* knockout strain may be attributed to biological autoluminescence, a phenomenon that has been found to be directly related to oxidative stress (Bereta et al. 2021). Together, these experiments developed an advanced modular assembly strategy, including selection, reporters, and an integration site for *X. dendrorhous* CBS 6398.

### Gene expression analysis for part derivation and elucidating photobiology

Of the current available promoters for *X. dendrorhous*, only P_adh4_ and P_act_ have high strength (Hara et al. 2014). Therefore, there is a need for more strong promoters. Furthermore, previous work has shown that astaxanthin metabolism is induced by oxidative stress and light (Zhang et al. 2019; Tropea et al. 2013; Stachowiak et al. 2013; Schroeder et al. 1995; Li et al. 2023; Liu et al. 2006; Kim et al. 2005; Vázquez et al. 2001), and Huang *et al*. has discovered a homolog of the white-collar 2 transcription factor in *X. dendrorhous* (Huang et al. 2022), hinting that *X. dendrorhous* may possess stress and light-inducible promoters. Transcriptomics, and particularly differential gene expression (DGE) analysis, is one way that both strong constitutive and regulated promoters could be identified.

Yet, it is not known that the response observed in astaxanthin biosynthesis is indeed transcriptionally mediated. Thus, before conducting transcriptomics, we used the genome sequence to design quantitative polymerase chain reaction (qPCR) probes to target genes in the terpenoid (*MVK*, *PMVK*, *MVD*, *IDI*, *FPS*, and *crtE*) and carotenogenic (*crtYB*, *crtI*, and *crtS*) pathways. We compared the expression of these genes in the dark and under ultraviolet light exposure (Methods). The log2 fold change (light/dark) of each gene in the pathway is depicted in Figure 2. Upstream genes *MVK*, *PMVK*, *MVD*, *IDI*, *FPS*, and *crtE* exhibited little to no change in expression between dark and ultraviolet conditions. However, expression of *crtYB*, *crtI*, and *crtS* was upregulated. Notably, the *crtI* gene showed the highest log2 fold change. This indicates that upstream terpenoid synthesis to the diterpenoid intermediate, geranylgeranyl pyrophosphate, is unaffected by light. These results are the first to demonstrate that light, particularly ultraviolet, transcriptionally upregulates carotenogenesis, not the upstream terpenoid pathway, in *X. dendrorhous*.

**Figure 2.**
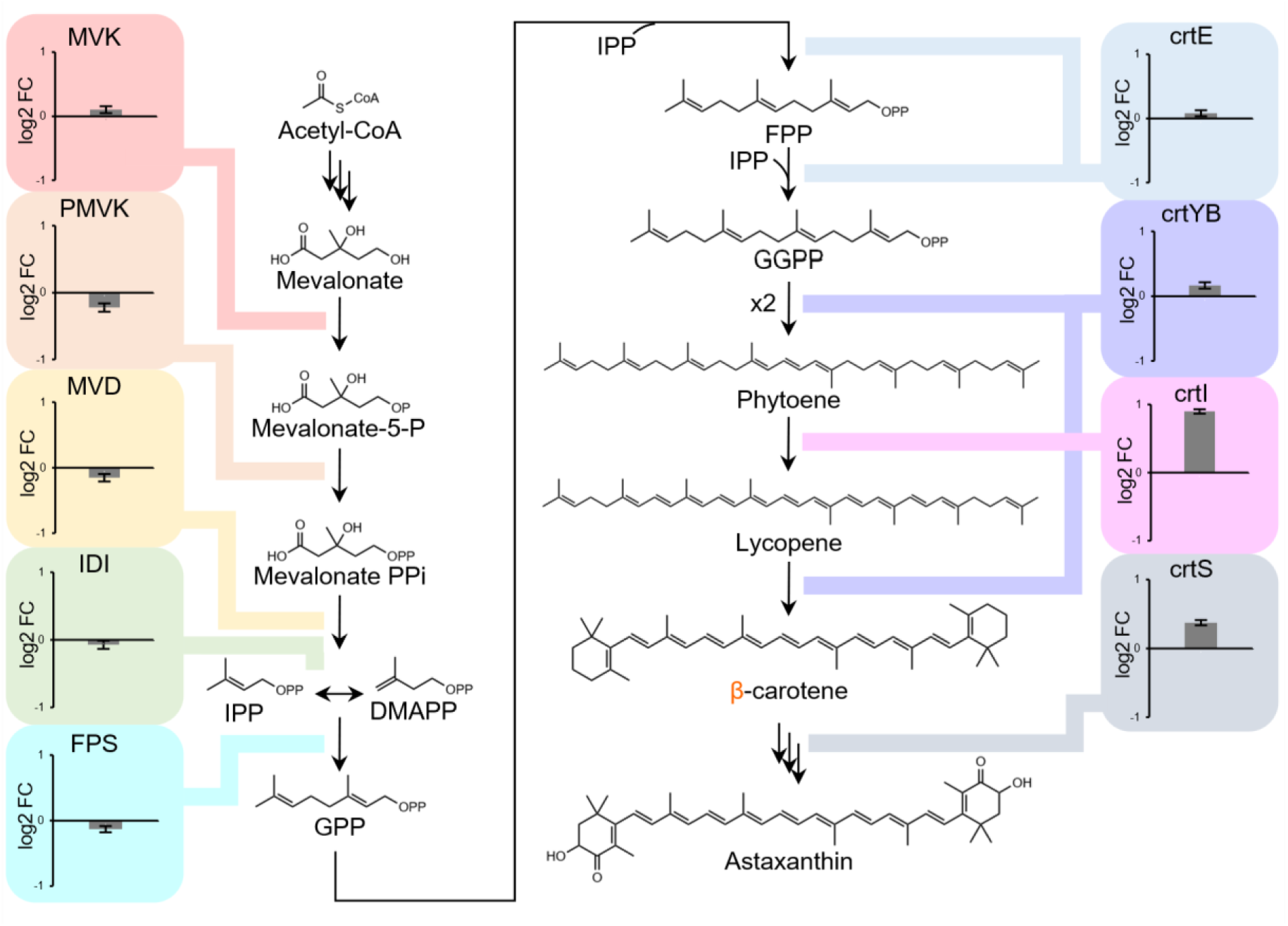
Transcriptional response of terpenoid and carotenoid biosynthesis in response to ultraviolet light exposure. Differential expression between dark and ultraviolet as measured by the log2 fold change (log2FC) of RT-qPCR ΔΔCt values (Methods). Upstream mevalonate and terpenoid genes are not affected by ultraviolet, but downstream carotenoid biosynthetic genes *crtI* and *crtS* are upregulated, *crtI* most strongly. *Abbreviations:* MVK – mevalonate kinase, PMVK – phosphomevalonate kinase, MVD - diphosphomevalonate decarboxylase, IDI - isopentenyl-diphosphate isomerase, FPS - farnesyl diphosphate synthase, crtE - geranylgeranyl pyrophosphate synthase, crtYB - bifunctional lycopene cyclase/phytoene synthase, crtI – phytoene desaturase, crtS - cytochrome-P450 hydroxylase.

Once we confirmed there was a transcription-mediated response to light, we designed a transcriptomics experiment to quantify differential gene expression across dark, ultraviolet, blue, green, and red light, with hydrogen peroxide oxidative stress as a control (Figure 3a). To perform the experiment, we constructed a Light Plate Apparatus (LPA) to culture *X. dendrorhous* under a variety of conditions (Gerhardt et al. 2016). Sequencing of the samples was done by the Department of Energy Joint Genome Institute, through their Community Science Program, and log 2 fold change in Fragments Per Kilobase of transcript per Million (FPKM) was calculated relative to gene expression in the dark using DESeq2 (Methods).

**Figure 3.**
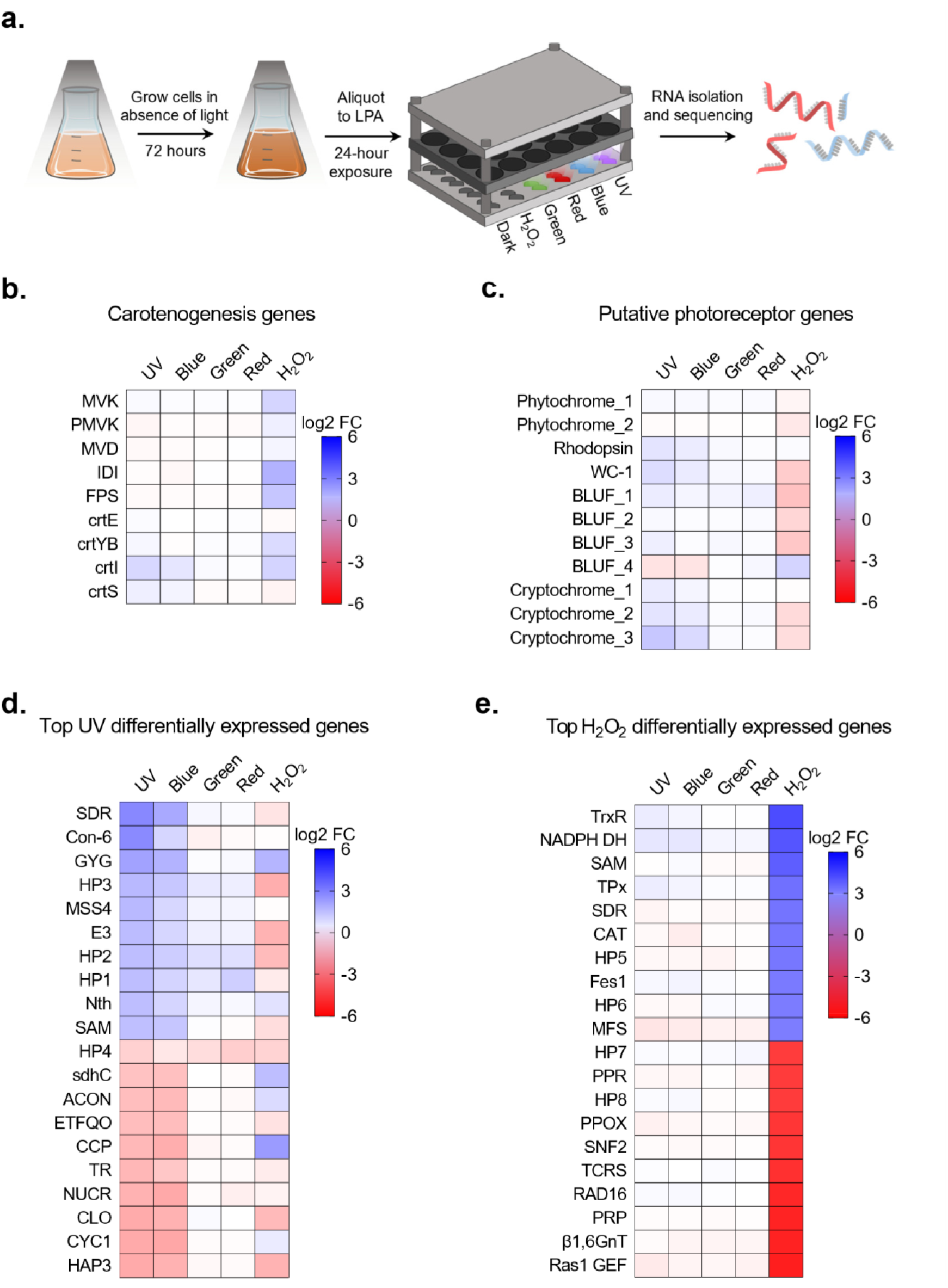
Transcriptomics and differential gene expression (DGE) of *X. dendrorhous* CBS 6398 under light exposure and oxidative stress. Blue represents induction and red represents repression. Color intensities are equally scaled across charts. **(a)** Depiction of the experimental workflow for obtaining RNA from *X. dendrorhous* cultured in the light plate apparatus (LPA) under light and hydrogen peroxide stress conditions. **(b)** Regulation of mevalonate and carotenogenesis genes by light and H_2_O_2_. Expression of *crtI* and *crtS* is considerably induced by ultraviolet light, and *crtI* is also induced by oxidative stress. **(c)** Transcriptional response of putative photoreceptor homologs identified by BLAST. The numbers for genes within each class are arbitrarily assigned. **(d)** Top ten ultraviolet upregulated and downregulated genes. *Abbreviations:* HP1-4 – hypothetical proteins, SDR – short chain dehydrogenase, Con-6 – conitiation-specific protein 6, GYG – glycogenin glucosyltransferase, MSS4 – phosphatidylinitosol-4-phosphate 5-kinase, E3 – E3 ubiquitin ligase, Nth – endonuclease III, SAM – S-adenosyl-L-methionine-dependent methyltransferase, sdhC – succinate dehydrogenase cytochrome b560 large subunit, ACON – aconitate hydratase, ETFQO – electron-transferring-flavoprotein dehydrogenase, CCP – cytochrome c peroxidase, TR – unclassified transcriptional regulator, NUCR – unclassified endonuclease, CLO – caleosin, CYC1 – cytochrome c, HAP3 – transcriptional activator heme activator protein. **(e)** Top ten H_2_O_2_ induced and repressed genes. *Abbreviations:* HP5-8 – hypothetical proteins, TrxR – thioredoxin reductase, NADPH DH – NADPH dehydrogenase, SAM – S-adenosyl-L-methionine-dependent methyltransferase, TPx – thioredoxin reductase, SDR – short chain dehydrogenase, CAT – catalase, Fes1 – Fes1-domain containing protein, MFS – major facilitator superfamily permease, PPR – pentatricopeptide repeat protein, PPOX – protoporphyrinogen oxidase, SNF2 – SNF2 family DNA-dependent ATPase, TCRS – two-component sensor, RAD16 – DNA repair protein, PRP – pre-mRNA-processing-splicing factor, β1,6GnT – beta-1,6-N-acetylglucosaminyltransferase, Ras1 GEF – Ras1 guanine nucleotide exchange factor.

### Transcriptomics elucidates *X. dendrorhous* photobiology

Before deriving promoters, we first analyzed the DGE data to understand the *X. dendrorhous* light response. We examined the terpenoid and carotenoid pathway (Figure 3b), as well as putative photoreceptor genes (Figure 3c) and the top differentially expressed genes in ultraviolet and oxidative stress (Figure 3d and 3e, respectively).

The terpenoid pathway data corroborated our previous evidence that exposure to ultraviolet light leads to upregulated expression of *crtI* with a log2 fold-change of 0.90. This is below the typically accepted significant log2 fold-change of 1.0, yet the other genes in terpenoid and carotenoid metabolism had no significant changes even at a low threshold of log2 fold-change 0.5. In contrast, oxidative stress upregulated genes in both pathways (*MVK*, *PMVK*, *IDI*, *FPS*, *crtYB*, and *crtI*), with *IDI* having the greatest increase (log2 fold-change of 1.79). This finding is significant because *IDI* is a known bottleneck for the mevalonate pathway in other organisms (Shin et al. 2022). Therefore, a deregulated *IDI* could be a target for increasing flux through the terpenoid pathway.

The light response systems in *X. dendrorhous* are relatively unknown, save white collar (Huang et al. 2022). However, systems such as cryptochrome, BLUF-domain, rhodopsin, and photolyase are known in other fungi, including a distantly related basidiomycete *Ustilago maydis* (Yu & Fischer 2019; Brych et al. 2016; Brych et al. 2021). Therefore, we searched the *X. dendrorhous* CBS 6938 genome with BLAST for putative fungal photoreceptor proteins within the BLUF, rhodopsin, white collar, and cryptochrome families (Figure 3c). Photoreceptor proteins predicted to be in the same family were differentiated with arbitrarily assigned numbers. We found that the putative cryptochromes increased transcription most dramatically in ultraviolet and blue light exposure. Additionally, putative white collar and rhodopsin genes increased expression. To our knowledge, this is the first evidence of cryptochrome, rhodopsin, and BLUF-domain light response systems in *X. dendrorhous*. The oxidative stress condition corroborated that these genes were responding specifically to light because most putative photoreceptors were downregulated or unaffected by hydrogen peroxide.

Finally, we filtered the whole genome transcriptomic data for the twenty genes most up- or downregulated by ultraviolet light. Of these twenty, ten upregulated and six downregulated genes had log2 fold-changes of 1.50 or greater (Figure 3d). Upregulated genes include short-chain dehydrogenase (SDR) which is involved in the induction of light-regulated pathways (SDR) (Bruchez et al. 1996), MSS4-like protein (MSS4) which is involved in cell survival through cell cycle regulation and actin cytoskeleton formation (Desrivières et al. 1998), and endonuclease III (Nth) which is directly involved in response to ultraviolet stress through DNA repair (Serafini & Schellhorn 1999). These functions collectively correspond to an expected stress response from ultraviolet exposure (Sinha & Häder 2002). Another upregulated gene, conidiation-specific protein 6 (Con-6), is involved in the reproductive cycle and is also upregulated after blue-light exposure in *Neurospora crassa* (Olmedo et al. 2010). Unexpectedly, ultraviolet light upregulates expression of glycogenin glucosyltransferase (GYG) (Goldraji et al. 1996, Smythe & Cohen 1991), an enzyme that initiates glycogen nucleation, and E3 ubiquitin ligase (E3), which is involved in growth and protein turnover (Cao & Xue 2021). Also of note, four of the ten upregulated genes were predicted to result in hypothetical proteins, indicating that our understanding of light response mechanisms is incomplete. Several downregulated genes in ultraviolet are involved in the cytochrome c system, including cytochrome c (CYC1) itself, cytochrome c peroxidase (CCP) (Pelletier & Kraut 1992), and the heme activator protein (HAP3) involved in cytochrome c regulation (Hahn & Guarente 1988). Others are mitochondrial electron-transferring-flavoprotein dehydrogenase (ETFQO) which is involved in the electron transport chain (Toplak et al. 2019), and caleosin (CLO) which is usually upregulated during stress (Rahman et al. 2018), succinate dehydrogenase cytochrome b560 (SdhC) (Abraham et al. 1994), and aconitate hydratase (ACON) which is an indicator of oxidative stress (Rakhmanova et al. 2023). Two further genes are unclear but labeled as a transcriptional regulator (TR) and endonuclease (NUCR). We then identified the ten most upregulated and ten most downregulated genes in the hydrogen peroxide exposure condition. Many of these genes are expected, and are different from the genes in the light conditions, supporting the idea that light and oxidative stresses elicit separate responses (Figure 3e).

To more clearly analyze the overall metabolic response to ultraviolet, genes were clustered into functional groups using the ERGO™ Bioinformatics software from Igenbio. Upregulation of functional groups is depicted in Figure 4a, along with their regulation pattern under oxidative stress. Generally, DNA repair and amino acid metabolic functions were upregulated. Interestingly, aromatic amino acid degradation and modification were upregulated by both ultraviolet light and hydrogen peroxide, but amino acid uptake was only upregulated in ultraviolet. Furthermore, electron transport, glycolysis, gluconeogenesis, and carbohydrate metabolism were downregulated (Figure 4b). These results support the general trends observed in the analysis of individual genes. We synthesized these observations into a qualitative description of the *X. dendrorhous* CBS 6938 metabolic shift in response to ultraviolet (Figure 4c). With this view, the transcriptional response indicates that *X. dendrorhous* likely redirects flux from central carbon metabolism to carotenoids and aromatic amino acids upon ultraviolet exposure. Aromatic amino acids absorb ultraviolet and are also precursors to other ultraviolet light absorbing molecules like melanin and mycosporine compounds (Llewellyn et al. 2020; Gilchrest et al. 1996; Bhatia et al. 2011). The activation of aromatic amino acid pathways in response to ultraviolet has not been previously observed in basidiomycetes. These results indicate that the response to ultraviolet in *X. dendrorhous* is more extensive and complex than acting on carotenogenesis alone. Furthermore, the significant fraction of hypothetical proteins hints at additional unknown mechanisms that also contribute to the response.

**Figure 4.**
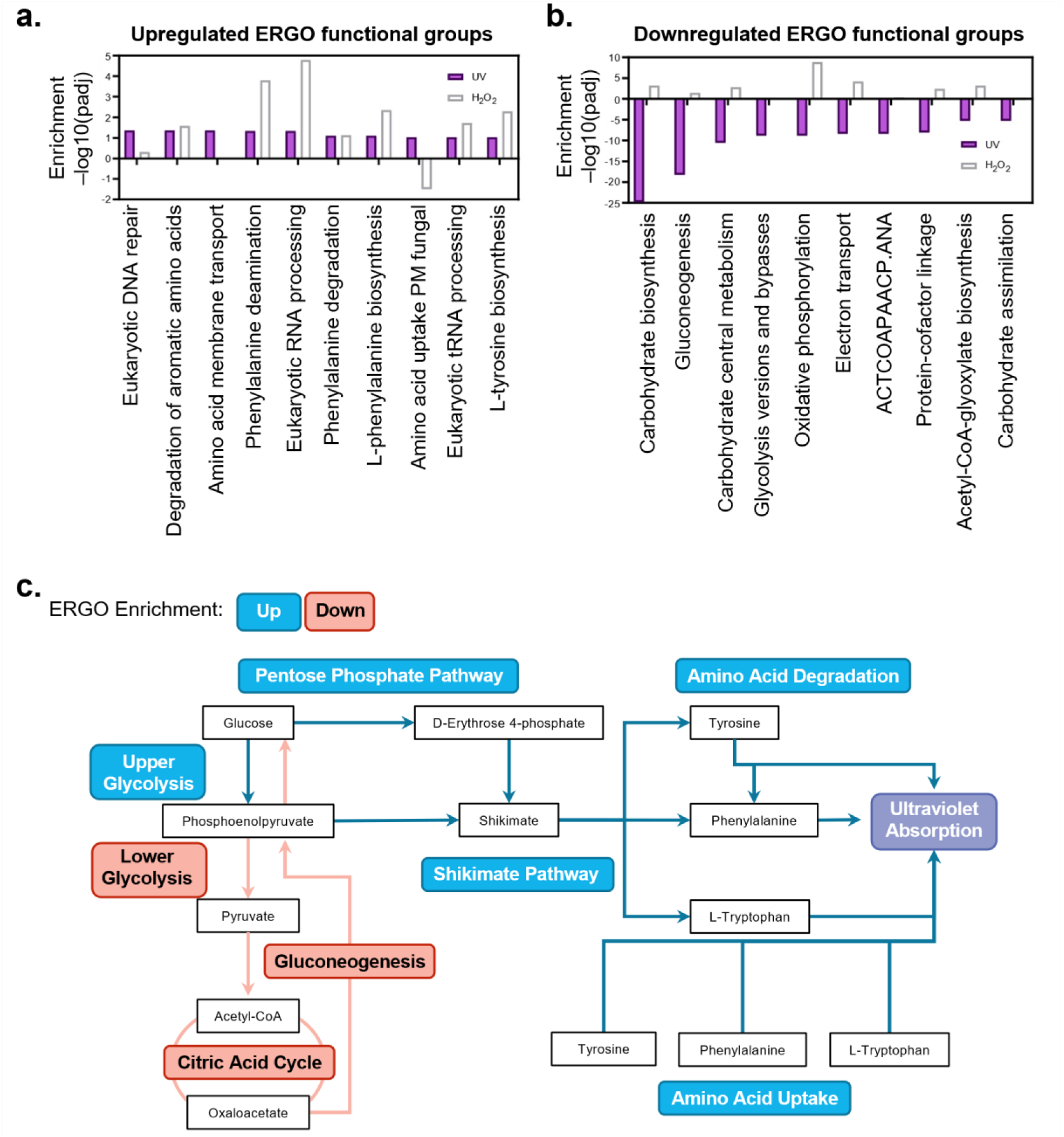
Transcriptomic analyses clustered by ERGO functional groups. **(a)** Overall upregulation of metabolic processes by ultraviolet light or H_2_O_2_. **(b)** Overall downregulation of metabolic processes by ultraviolet light or H_2_O_2_. **(c)** Summary of metabolic processes regulated by ultraviolet light. Generally, metabolic flux is being drawn from central carbon metabolism to synthesis of aromatic amino acids.

### Transcriptomics-driven part selection allows simultaneous derivation of constitutive and regulated genetic parts

With initial analysis complete, we then sought to leverage the DGE data to derive gene expression parts as originally intended. Since the data covered a variety of conditions, we reasoned that it should be possible to derive putative strong constitutive and regulated promoters. We identified strong constitutive promoters by sorting each transcriptomics dataset by fragments per kilobase of exon per million mapped fragments (FPKM) count, choosing the genes that had the highest and consistent FPKM counts across conditions (Figure 5a). Notably, the known strong *ADH* promoter is among these, yet no other central carbon metabolism promoters are represented. This result contests the convention of simply selecting central carbon metabolism promoters as genetic parts for yeasts. We also selected promoters from the genes in Figure 3d that had log2 fold-changes of 1.50 or greater when exposed to ultraviolet light (Figure 5b and 5c, respectively). We used the resequenced genome to obtain the promoter regions upstream of each of these 26 promoters, as defined by a JGI promoter analysis tool. These putative promoters were then cloned into our modular cloning system to drive expression of the luciferase reporter and integrated into the genome (Figure 5d).

**Figure 5.**
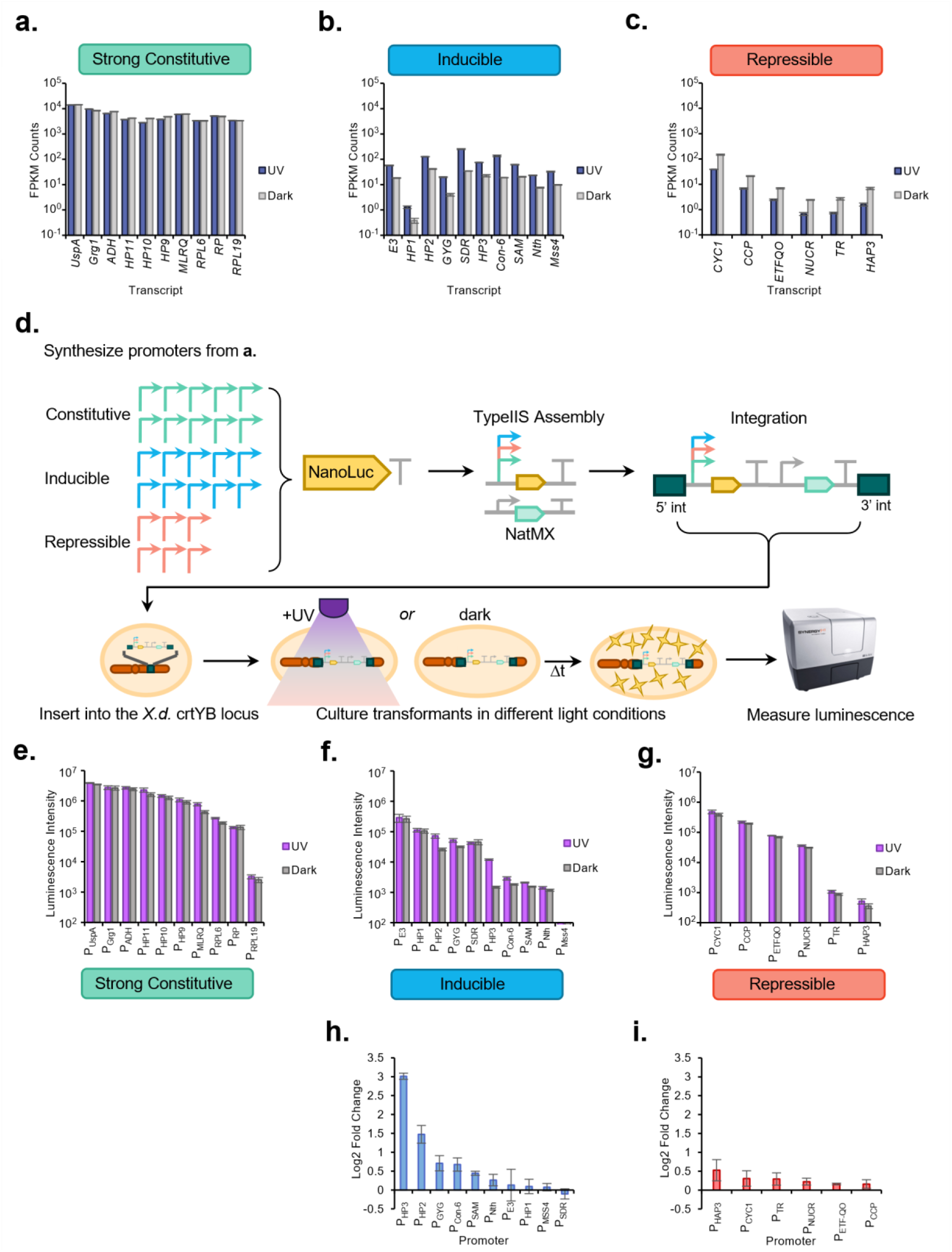
Deriving constitutive and regulated promoters from the transcriptome. **(a)** FPKM counts of the ten strongest constitutive promoters. *Abbreviations:* HP9-11 – hypothetical proteins, UspA – universal stress protein A, Grg1 – glucose-repressible protein 2, ADH – alcohol dehydrogenase, MLRQ – NADH-ubiquinone reductase complex 1 MLRQ subunit, RPL6 – ribosomal protein L6P, RP – ribosomal protein, RPL19 – ribosomal protein L19E. **(b)** FPKM counts of the top 10 inducible promoters, also shown in Figure 3d. **(c)** FPKM counts of the top 6 repressible promoters, also shown in Figure 3d. **(d)** Hierarchical DNA assembly and expression characterization workflow for each of the 26 promoters. **(e)** Expression strength of the strong constitutive promoters as measured by Nanoluc luminescence for recombinant *X. dendrorhous* strains grown in the dark or exposed to ultraviolet light. **(f)** Expression strength of the ultraviolet inducible promoters as measured by Nanoluc luminescence for recombinant *X. dendrorhous* strains grown in the dark or exposed to ultraviolet light. **(g)** Expression strength of the ultraviolet repressible promoters as measured by Nanoluc luminescence for recombinant *X. dendrorhous* strains grown in the dark or exposed to ultraviolet light. **(h)** Log2 fold change of luminescence intensity for the inducible promoters. **(i)** Log2 fold change of luminescence intensity for the repressible promoters.

Then, expression of each construct via a luciferase assay in the dark and under ultraviolet exposure was measured (Methods). As Figure 5e shows, the constitutive promoters result in high luminescence that does not change between the dark and the light. Many of the putative activated promoters showed increased luminescence in response to ultraviolet (Figure 5f). However, most of these increases were not large. The two strongest inducible promoters were found to be hypothetical proteins HP3 and HP2 with log2 fold-changes of 3.01 and 1.47, respectively (Figure 5h). The putative repressed promoters did not show any significant repression (Figure 5g and 5i). We reasoned that the long half-life of luciferase was confounding measurement of repression. Thus, we designed a qPCR experiment, and the results indicated no change in transcript levels (data not shown). Therefore, it seems likely that the promoter sequences selected do not contain the regulatory elements, or there could be a variety of context-dependent epigenetic factors.

Taken together, our results show that transcriptomics driven part discovery can consistently derive strong constitutive promoters, but inconsistently derive regulated promoters due to a variety of possible factors. Even so, we were able to discover ten strong promoters as well as two promoters that can be activated by ultraviolet, one of which that has 9-fold induction upon exposure to ultraviolet light. Thus, our part discovery efforts generated a collection of constitutive and regulated promoters that are compatible with our modular cloning system for *X. dendrorhous* CBS 6938.

## DISCUSSION

In this study, a promising nonconventional yeast was subjected to a streamlined set of genomics and genetics experiments that transformed it into a host for advanced genetic engineering and synthetic biology. This was done by obtaining a high-quality genome, developing advanced modular genetic tools, measuring gene regulation, and deriving a collection of genetic from transcriptomic data. Now, it is possible to interrogate the unique metabolism and photobiology of *X. dendrorhous* in a similar manner to other nonconventional yeasts.

Our analysis of *X. dendrorhous* photobiology revealed novel insights into the biological mechanisms of basidiomycete light response. We found evidence of multiple different fungal light response systems, including the first indication of cryptochromes, rhodopsins, and BLUF-domains in *X. dendrorhous*. Although previous studies have indicated that light increases astaxanthin production (An et al. 1989; Ojima et al. 2006; Leiva et al. 2015; Gutiérrez et al. 2015), we determined for the first time that light transcriptionally regulates the carotenoid, or tetraterpenoid, pathway but not the upstream terpenoid pathway. The key node, *IDI*, is also regulated by the oxidative stress response. We learned that *X. dendrorhous* engages a variety of survival responses in response to ultraviolet, including activation of DNA repair and reproduction pathways and repression of central carbon metabolism and mitochondrial respiration. We also observed aromatic amino acid biosynthesis is upregulated in addition to tetraterpenoid biosynthesis. Taken together, these data paint a picture of an extensive and coordinated response to light. This complex system is likely key to survival of *X. dendrorhous* in its ecological niche of fallen trees at high altitudes (Libkind et al. 2007), which would include extended periods of intense ultraviolet exposure.

It is interesting to note that our genome integration approach results in efficient on-target insertion of constructs via homologous recombination (HR). Our findings support the results of others that *X. dendrorhous* CBS 6938 has an active HR pathway (Hara et al. 2014), which is incredibly beneficial for future strain engineering efforts. This challenges the dogma that nonconventional yeasts favor the non-homologous end joining (NHEJ) mechanism, and argues that other yeasts besides *S. cerevisiae* may favor HR (Schwartz et al. 2017; Schwarzhans et al. 2016).

Leveraging transcriptomics allowed us to derive constitutive and regulated gene expression parts. Only one of the ten strongest promoters was from central carbon metabolism, challenging the practice of simply deriving promoters from central carbon metabolism for nonconventional yeasts. Transcriptomics has been previously used to identify regulated promoters in other yeasts (Gasser et al. 2015; Brink et al. 2023; Zahrl et al. 2017). This work builds on this prior evidence, adding a modular cloning standard to create an -omics-to-parts workflow that can yield needed constitutive and inducible promoters for construction of genetic devices. This -omics-to-parts workflow can be generalized to derive genetic parts collections for nonconventional yeasts.

Development of novel production platform hosts requires an understanding of genome composition and genetic regulation, which also is essential for genetic tool and genetic part development. This work demonstrates how an integrated genomics and genetics approach can simultaneously deliver new biological understanding and onboard a nonconventional organism, particularly a nonconventional yeast with no closely related model organism. This integrated approach, consisting of a small number of experiments, promises to accelerate organism onboarding efforts.

## MATERIAL AND METHODS

### Strains and media

The *X. dendrorhous* type strain CBS 6938 (ATCC 96594) was cultured at 21 °C in YPD (YEP, Sunrise Science, 1877-1KG plus 20 g/L glucose, Alfa Aesar, A16828) media for all experiments. For all light plate apparatus experiments, the *X. dendrorhous* seed cultures were shielded from any external light by wrapping the flasks with aluminum foil. *X. dendrorhou*s transformants were selected using YPD with 30 mg/L nourseothricin (Jena Bioscience, AB-101-10ML). Solid media plates were made using 20 g/L agar (Sunrise Science, 1910-1KG). Golden gate (Type-IIS) cloning was conducted using chemically competent *Escherichia coli* DH5α cells (NEB, C2987H). Construction of destination vectors with the toxic selection marker ccdB was done with chemically competent One Shot ccdB Survival 2 T1R *E. coli* cells (Invitrogen, A10460). Both *E. coli* strains were grown in 25 g/L LB Miller broth (Fisher Scientific, BP1426-2) at 37°C. Antibiotic selection was carried out using 100 mg/L carbenicillin (Alfa Aesar, J61949), 25 mg/L chloramphenicol (Alfa Aesar, B20841), and 50 mg/L kanamycin (Alfa Aesar, J61272) for level 0, level 1, and level 2 assemblies, respectively.

### gDNA isolation and sequencing

Isolation of gDNA from *X. dendrorhous* and next-generation sequencing was performed as described by Collins et al. 2021.

### Construction of a light plate apparatus for various light wavelengths

The light plate apparatus (LPA) was assembled following a user’s manual published by the Tabor lab at Rice University (Gerhardt et al. 2016). The LPA is an instrument capable of shining two individual LED lights on cell cultures in a 24-well plate. It is comprised of a 3D-printed shell surrounding a soldering board with 48 LED light sockets oriented below a 24-well plate with a clear bottom (AWLS-324042, Arctic White). The LPA allows each culture to be exposed to two unique LED lights without disrupting neighboring cultures.

### RNA isolation from the light plate apparatus

LPA cultures were started in 2 mL of YPD media at OD_600_ = 1 and shaken for 4 hours at 200 rpm while exposed to one red (λ = 660 nm), green (λ = 565 nm), blue (λ = 470 nm), white (visible light spectrum), or UV LED light (λ = 400 nm), or no LED light. For the transcriptome sequencing experiments, the hydrogen peroxide condition replaced the white light condition. The hydrogen peroxide condition consisted of YPD media with hydrogen peroxide at a concentration of 10 mM and had no LED light. 1 mL of *X. dendrorhous* culture was taken forward for RNA isolation, which used a cell homogenization and TRIzol-based method. *X. dendrorhous* cells were first resuspended in 200 μL of cell lysis buffer (0.5 M sodium acetate, 5% SDS, 1 mM EDTA) and transferred to pre-filled tubes with 400-micron zirconium beads (OPS diagnostics, PFMB 400-100-34). The cells were homogenized using a FastPrep-24 machine from MP Biomedical for two cycles at 6 m/s for 30 seconds each, with a 1-minute incubation on ice between each cycle. 800 μL of Trizol reagent (Invitrogen, 15596026) was then added to the cell lysate and incubated on ice for 10 minutes, followed by two more cycles on the FastPrep machine. On ice, 100 μL of 1– bromo–3–chloropropane (Sigma Aldrich, B9673-200ML) was added to the cell lysate, inverted to mix, and incubated for 5 minutes. The tubes were spun down at 4°C for 14,000 x g for 15 minutes. 400 μL of supernatant was transferred to 400 μL of 100% ethanol. The RNA was purified using the Monarch Total RNA Miniprep Kit (NEB, T2010S) following the supplier’s instructions.

### qRT-PCR evaluation of carotenogenesis pathway activity in response to light exposure

RNA samples were converted to cDNA using the Protoscript II First Strand cDNA Synthetsis Kit (NEB, E6560S) following the supplier’s standard protocol instructions. RT-qPCR was then performed using IDT’s PrimeTime Gene Expression Master Mix (Integrated DNA Technologies Inc., Skokie, Illinois, 1055772) and PrimeTime qPCR pre-mixed assay with probes and primers. Sequences can be found in Table S2. Probes targeting genes-of-interest were labeled with FAM fluorophores and probes targeting the actin housekeeping gene were labeled with HEX fluorophores. qRT-PCR was performed in triplicate. Primers were designed on Benchling (https://benchling.com/) and optimal probe placement was determined using PrimerQuest (https://www.idtdna.com/PrimerQuest). The QuantStudio 6 Flex system was used, and instructions provided by IDT for the standard cycling protocol were followed.

### RNA transcriptome sequencing and differential gene expression analysis

cDNA preparation and sequencing was performed per JGI’s standard workflow described in Girgoriev et al (Kim et al. 2005). Raw FASTQ file reads were filtered and trimmed using the Joint Genome Institute (JGI) QC pipeline (Clum et al. 2021). Filtered reads from each library were aligned to the reference genome using HISAT version 0.1.4-beta (Kim et al. 2015). featureCounts was used to generate the raw gene counts file using gff3 annotations (Liao et al. 2014). Raw gene counts were used to evaluate the level of correlation between biological replicates using Pearson’s correlation and determine which replicates would be used in the DGE analysis. DESeq2 (version 1.10.0) was subsequently used to determine which genes were differentially expressed between pairs of conditions (Love et al. 2014). The parameter used to call a gene differentially expressed between conditions was a p-value < 0.05. Three biological replicates were performed for each treatment. Promoter names were determined by comparing gene annotations across ERGO’s gene ontology, Interpro, JGI predictions, and BLAST results, as shown in Table S4.

### Gene set enrichment analysis

Annotated *X. dendrorhous* CBS 6938 genome and gene count table from mapped transcriptomic reads were uploaded to ERGO 2.04 (Igenbio, inc) (Overbeek et al. 2003). Genes were functionally enriched according to ERGO groups using GAGE5 (Luo et al. 2009).

### Polymerase Chain Reaction (PCR)

All PCRs were done using Q5 2X Master Mix (NEB, M0492L). Primers were designed on Benchling (https://benchling.com/) and the NEB Tm Calculator (https://tmcalculator.neb.com/). All primers were ordered from IDT. PCR reactions closely followed NEB instructions. Briefly, reactions were done in 50 μL total volume; 25 μL Q5 Master Mix, 2.5 μL of each primer, X μL of template DNA (1 ng plasmid DNA or 100 ng genomic DNA), and 20-X μL nuclease free water (VWR 02-0201-0500). Reactions were run on a thermocycler with the following manufacturer instructions.

### Gibson cloning

Gibson assembly was used to construct destination vectors, which were designed for TypeIIS-based cloning. Reactions closely followed instructions from the NEBuilder HiFi DNA Assembly Master Mix (NEB, E2621S). Fragments were amplified with PCR and designed to have 20-30 bp overlaps. The DpnI enzyme was used to digest template plasmid or *X. dendrorhous* genomic DNA according to the manufacturer’s instructions (NEB, R0176S). The amplified fragments were purified using the Zymo Clean & Concentrator Kit (Zymo, D4005), followed by dilution to 0.2 pmols for 2-3 fragments or 0.5 pmols for 4 or more fragments. The fragments, 10 μL of HiFi master mix, and nuclease free water were combined in a PCR tube for a 20 μL total reaction volume. The mixture was run on a thermocycler at 50°C for 60 minutes, with a hold at 10°C.

### TypeIIS Cloning and Parts

A modular, hierarchical TypeIIS cloning scheme was used construct genetic designs for integration into *X. dendrorhous*. This scheme consisted of three levels; transcriptional parts (level 0), transcription units (level 1), and integrative pathways (level 2). Cloning reactions were based on the enzymes BbsI (Thermo Scientific, ER1011) or BsaI (NEB, R3733). Cloning reactions consisted of 1 μL of N parts, 1 μL of BsaI or BbsI, 1 μ L of 10X Ligase Buffer, 0.4 μL of T4 DNA ligase (Promega, M1794), and 7.9 - N μL of nuclease free water, where N is the number of parts. Reactions were run on a thermocycler at 37°C for 5 hours, 50 °C for 15 minutes, 80 °C for 20 minutes, and a hold at 10°C. All genetic parts used in this study can be found in Supplementary Table 1. All cloning vector maps can be found in Figure S2.

### Minimum Inhibitory Concentration Assay

*X. dendrorhous* cultures were diluted with YPD to an OD_600_ = 1.0 and 100 μL was pipetted into columns 2-11 of a 96-well plate (Corning, 3596). Column 11 served as a positive control of only *X. dendrorhous* culture. A negative control of 100 μL YPD was pipetted into column 12. The antifungals hygromycin (ThermoFisher, 10687010), zeocin (Jena Bioscience, AB-103S), geneticin (ThermoFisher, 10131-035), and nourseothricin (Jena Bioscience, AB-101L) were chosen for the experiment. Stocks of each antifungal were made in concentrations of 5120 μg/mL and 7680 μg/mL. Five biological replicates of antifungal concentration were made by pipetting 200 μL of an antifungal into three wells of column 1. A multichannel pipette was used to perform a serial dilution to column 10. At column 10, 100 μL of the culture and antibiotic mixture was pipetted and discarded, leaving 100 μL of mixture in every well. The 96-well plates were incubated at 21°C for 3 days. The OD_600_ was measured on a BioTek Synergy H1 plate reader with column 12 as a blank. Survival percentage was calculated by the following equation: 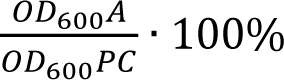 where OD_600_A represents the average OD_600_ associated with a certain antibiotic concentration, and OD_600_PC represents the average OD_600_ for the positive controls in column 11. Outliers were calculated using an interquartile range and were excluded in survival percent calculations. Outliers were distributed nearly evenly across all four antibiotics treatments with 16 in the hygromycin data, 13 in the nourseothricin data, 15 in the zeocin data, and 20 in the geneticin data. Outliers were calculated and removed due to imperfect measurements of plate readers with heavy yeast cells, such as *X. dendrorhous*.

### X. dendrorhous transformation

Integrative pathway DNA for transformation was excised from level 2 vectors with a BsaI digestion. This digestion was comprised of 40 μL plasmid DNA, 5 μL 10X CutSmart Buffer, 2 μL BsaI, and 3 μL nuclease free water. The reaction was run at 37°C for 10 hours, 80°C for 20 minutes, and a hold at 10°C. The resulting DNA fragments were then purified with the Zymo Clean & Concentrator Kit and were ready for transformation. *X. dendrorhous* transformations were based on an electroporation method described by Visser et. al. 2003. *X. dendrorhous* was first streaked on a YPD agar plate and grown at 21°C for approximately 2 days. A single colony was used to inoculate 50 mL YPD media in a 125 mL Erlenmeyer flask. The cells were shaken at 21°C and 200 rpm for another 2 days. These cells were then used to inoculate 200 mL of YPD in a 1 L Erlenmeyer flask at OD_600_ = 0.02. The cells were grown at 21°C and 200 rpm until reaching OD_600_ = 1.2 (approximately 20 hours). From here, the cells were pelleted at 1500 x g for 5 min and resuspended in 25 mL of freshly made 50 mM potassium phosphate buffer (pH=7.0, Sigma-Aldrich, P8281-100G & P9791-100G) with 25 mM diothiothreitol (Acros Organics, 426380500). The cells were incubated at room temperature for 15 minutes, and then pelleted at 4°C for 5 minutes at 1500 x g. The cells were then washed with 25 mL of ice cold STM buffer (pH = 7.5, 270 mM sucrose, Millipore Sigma, 1.07651.5000, 10 mM Tris HCl, Alfa Aesar, J67233, 1 mM MgCl_2_, VWR, E525-100ML) and spun down again at 4°C for 5 minutes at 1500 x g. This wash step was repeated and followed by resuspension of the pellet in 500 μL of STM buffer. Now electrocompetent, the cells were divided into 60 μL aliquots and kept on ice until electroporation. 10 μL of transforming DNA was mixed with a 60 μL aliquot and transferred to an ice cold 0.2 cm electroporation cuvette (ThermoFisher, 21-237-2). Electroporation was performed with a Gene Pulser electroporator (Bio-Rad, 1998.018.1, 1998.018.2, & 1998.018.3) at 0.8 kV, 1000 ohms, and 25 μF. Immediately following the electric pulse, 500 μL of ice cold YPD was added to the cuvette. This mixture was then transferred to 4 mL of YPD in a 14 mL Falcon tube and grown overnight on a rotating drum. The next morning, the cells were spun down for 5 minutes at 1500 x g, resuspended in 500 mL of filtered water, and spread onto selection plates. The plates were incubated at 21°C until colonies appeared, which typically occurred in 2-3 days. All strain modifications performed on CBS 6938 can be found in Table S3.

### Astaxanthin extraction from wild-type and *crtYB* knockouts

*X. dendrorhous* CBS 6938 wild-type and *crtYB* knockout strains were grown for 3 days in the dark at 21°C and 200 rpm. 1 mL of culture was taken forward and centrifuged at 10,850 x g for 10 minutes. The cell pellet was washed with Milli-Q filtered water twice before resuspension in 1 mL of acetone (SigmaAldrich, 34850-1L). Cell and acetone mixtures were poured into pre-filled tubes of 400-micron zirconium beads (OPS Diagnostics, PFMB 400-100-34), then transferred to snap-cap microcentrifuge tubes (Eppendorf, 022363743). The Bullet Blender Storm Pro tissue homogenizer (NextAdvance, BT24M) was used at 12 m/s for 5 minutes to mechanically lyse the cells. Afterward, tubes were centrifuged for 10,850 x g for 10 minutes. 200 μL of supernatant was pipetted into a nylon syringeless filter (Whatman, UN203NPENYL) and transferred to a 2 mL chromatography vial (Agilent, 5181-3376) with a glass vial insert (5183-2085) and crimp top cap (Agilent, 8010-0051).

### Ultra-high performance liquid chromatography detection of astaxanthin

UPLC analysis was performed using a modified method from Bohoyo-Gil et al. 2012. Modifications include an injection volume of 10 μL, usage of a Shimadzu Nexcol C18 1.8 μm column (Shimadzu, 220-91394-03), and analysis on the Nexera Series UPLC (Shimadzu; RF-20AXS, RID-20A, SCL-40, DGU-403, DGU-405, CTO-40C, SPD-M40, C-40 LPGE, LC-40D XS, SIL-40C XS).

### Bioluminescence detection assay

LPA cultures were started in 2 mL of YPD media at OD_600_ = 1 and shaken for 24 hours at 200 rpm. *X. dendrorhous* was exposed to either a UV (λ = 400 nm) or dark (no LED light) condition. Bioluminescence was detected using the Nano-Glo Luciferase Assay System (Promega, N1110). Assay buffer and assay reagent were mixed in a 50:1 ratio to create to assay solution. Aliquots of 50 μL of *X. dendrorhous* cells were mixed with 50 μL assay solution in a 96-well plate (Corning, 3904). Bioluminescence was measured on a BioTek Synergy H1 plate reader.

## AUTHOR CONTRIBUTIONS

EET, JHC, and EMY conceived of the study. EET and JHC performed all experiments and analysis save the sequencing and initial processing of RNA-seq data. CBM and GTN constructed the light plate apparatus (LPA). KM acquired microscope images of red yeast. AL, SM, and IG performed transcriptomic sequencing and initial processing of RNA-seq data. EET and EMY wrote the manuscript.

* Emma E. Tobin and Joseph H. Collins contributed equally.

## COMPLIANCE WITH ETHICAL STANDARDS

## DECLARATIONS

This article does not contain any studies with human participants or animals performed by any of the authors.

## DATA AVAILABILITY

The genome was deposited in GenBank under the BioProject PRJNA1053634. Each replicate was registered as an NIH BioSample by JGI and can be found under the accessions SAMN15720006, SAMN15720353, and SAMN15720008 (UV1-3), SAMN15719986, SAMN15719959, and SAMN15720031 (blue1-3), SAMN15720054, SAMN15720004, and SAMN15719987 (green1-3), SAMN15719960, SAMN15719963, and SAMN15720352 (red1-3), SAMN15719962, SAMN15720053, and SAMN15719985 (H_2_O_2_1-3), SAMN15720005, SAMN15720007, and SAMN15719961 (dark1-3). Additionally, all genomics and transcriptomics data can be found within the JGI genome portal under the JGI Project ID 1255824.

