## Supplementary Information for "Omics driven onboarding of the carotenoid producing red yeast *Xanthophyllomyces dendrorhous* CBS 6938"

† Equal contribution

### SUPPLEMENTARY TABLES

**Table S1:** Genetic parts used in this study.

| Name | Type | Sequence | Source | Reference |
|---|---|---|---|---|
| $P_{UspA}$ | Promoter | AGCCTTGTTGCCTCTCTGGTATCTGAATCAAAAAACAGAAGTAGGGATTTCCATC<br>AGGTCTGGGTTCGAGTGTCATTGCATGTTGTGTTTTCTCTTTGGATCTCTATAGA<br>ACGCTGGCCTATTCTTCTGTGATGTTTTTCTTCCCTTTCTTTTTCTTCTTTCTTG<br>ACTCTCTTCATGCTTCTTCTTTCCTCTACATCAATTTCCATAGCTTCCCCCTTTA<br>CTCTTTTTCCCCATCAACCCTTTGTGTCTTTGTTCGTCTTGCGTATTTAACCTTA<br>TCCGTCATCGTTCCAAAGGCTTGTTTCGAATGGAAGAAGAAAAACATTATAAAAT<br>TCGATCTTTCATCTTGCCTTTTCCCTGTTTTGGAACGAGGATAGTGTTTTCTCAA<br>GAGAATGACGTCTCTTGAGAGCAAGGGTGAATGACGCGATCAGAACGGACGAATA<br>CGAGTGATTGAGCACAAGATAAGCTAGACGAGGAGAAGAAGAAGAAAAAATCAAG<br>AATTCGTTGGAATCTTCGGAGTCTTTCTTGTTTCTTCCTGATCGCCAGGACGTTT<br>CTCGACCGAAACTCTACTTTCGAGCTGTCGGTACTCACCCCGGACTGGACCGAAC<br>TCGTCCATCTTATTCCGGACAGAGTCGGTCTGCGTTCGGTTACCTCACCCACTGA<br>CGCTCAGACTGCCGAACGTACTGACAGCTCGGCATGGCATGGCCAGAAAGTAGGG<br>AAGAACCTTACGTCGGGACTGTCCGGCTCCACCCGATGGGCTAGGATACATGGCT<br>AACTTGTTCCGTCGCCATCTCTTTGAGTGGGCTCATGTATGAGCTTGAACTGTAT<br>AAAATGCTGACCGAGTTGGTGTTTCTGCACTGAGAGGTTTCATCGTTTCTTCACC<br>CCATCAGACTTCCACCAAGCAACTCCTTTCTACACATCTACCCAACGCTAACAAT<br>CTACTCGATCTACTTACAATCAACATC | This study | |
| $P_{Grg1}$ | Promoter | ATTTGTTGTCGCTTGTATCGATTGCTAAAACGAACGAAATGATGAAAGCAAGACT<br>GTGAAAGTCTATATCCGTAGTTAGGGTTTAGGTGTTCTAGGCAACGATATGGCCA<br>TGGATGTTTATCTTCAGTTGGGCCAGCTTCTCGATGGTTCTAGACTGTGAGCGCT<br>TTCTGTTTGGTTGAGAGAGAAATCTAGTCAATGGTGAGGGAAGCGCCTCGTATGC<br>GGGTGTGTGTACGTGTACCTGTACGCGTATGTACGTGAGCGAGCGCGCTCGAGGT<br>ATATGCGTTCCGCGAACTGCGGAGATGTGACGTCACATCTCGTCACAGCTCATCA<br>CATGACGTCGTCCAACTTAGATGAGCGATCGATATGATGACGGCGTCTGTGCCTT<br>GTGCCTTGTTCCTGACCGGGCTCGACTGTCGAACGAGCGGACCAGAGATGCCGGC<br>GAGATGGGCTGAGCTGTGACGACACGGTGGGACACGTTGAACATTGACAGGGGTT<br>TTTTCCTCCTTCGTCTGCTATATAAGAAGGCCGGCTGGATGACATTGGGGTTCGC<br>TTCTTCTTCCCTCTCACACACAAAGCATTTCTCACTCTTCACTCTACTCTCAACC<br>AAACCAACCACCCCAAACAAACACTAATCTTTCACA | This study | |
| $P_{ADH}$ | | GGTAAAGAAAAGAAAGTCGACCGTATAAGAACGGAGGATCCAAGAGAAGGCGACA<br>TTCTGAAGCAACCAGACGAGGAGATAGATGACTATGGATTCACAGAGTGATAGGG<br>TCAACGAACTCGATTAGTCAGCTTGATAACGCGCAGTAGCCTTCACGCGCAGCAG<br>CCACGTCATTTCGGCCCGGCGCAAACGGATGCCGAGTGTTTTTTTCCGGGCCGTT<br>CGGAAGAGAGAGGAACAACGACGATGGCAGAGAAAACATCTTGGGATGGACAACT<br>CTTGCTGTTCACGTCAGAGGCTGAGGACCAAGGGCTGGGGATTGGGGTTTGAGAT<br>TGACTCTAGAGTGATAGTAATATCTGGAGATCAGAGCTGAGACTGATAAGACTGA<br>TGCTAACGAGCTGACGCTGATAGATTACCGGTGCCGCTGAAACTGCTATCCGGAC<br>TAACGAGGTGGAAGATGACGGAACTCTTTCTTCGCCGGCTGTTCCATCTTCTTCT<br>TCGTCTTTCTCTCATTGCCTCGTTTTTCTTCCCATCGAGGGGCTCCTCAGAAACG<br>TCCGGTTTCCGACATCTTCTGTCAATCTTTGTTCCAGTCTGTGGTAGGTGCTTGT<br>GTGCTGATCCTTACGGCCTCTGTTCTTCCAGGGGTTTCCCGTGGGAAATTTGTCC<br>GGGAAGGAATCCGATTGATTCCTGACAATGTCCTTTCTTTCGAGCGATAGGCAGA<br>TGACTGGTTTATTTAATGTTTCCCTTTTCCGTCTGCTCGCTCACGGAGGATCCAA<br>CGTTCTTTCTCTTCCTCCACTTCCGTATCGTCCGGGGCAGATCGTCATCACAGAA<br>CGGAGACTGAAATGAACCATATAAAGCAGGTGGTGTCTCGATCTCAATCTCACTT<br>TCCTTTCTTTCCCTCACCATACAAACATCCCATCCTATCAAGCCTCAGCATCACT<br>CCTAGCTTACTCTCACAGAGATCCTATTCAACCAACCAACCAACAACACATCATC<br>TGAACCCCACA | This study | |
| $P_{HP11}$ | Promoter | CTAGCGGAAGGAGTGGGAGGAAGGGGATGGGACAAGAAGAGAGAGAGGAAGGAAG<br>AGAAAGAAGAACGTTGAGTATAGACAAGATAGAGTCGACAGTGCGGCAAAGACGA<br>TGAAAATGAGCGAAGGATTTCGATTAGACGGAATCGGCCGGGAGCGGGTAAGCGA<br>GGAAGGCTCGTGCGTGGCGAGGTCACGTGTGTTGTTGCGTTTGACACGTAACCAT<br>GATCTGGCCGACACCCCAAAAGCAGTCCAAACGAGGCCCAAGGCTTGTGGTTGAT<br>GATGTCAGAACCTCTGAGATTAGATTAGATTGGCATCGAACAAATCATTCGTCAG<br>CTCATCGAACTCTATCGATGGACGATTCGGATGACATCATTTGGCCAGAGCCTAC<br>ACAAAGGGAAGCGTCGACGGATGGCGCGTCCAGCCTGCGGACCCACAGGAGGGAC<br>CGATGGGACGAGATGGGTGCGGGAGAGATCCAGGCAGCGCAGGCACGAAAGCAGG<br>CATAGTCAGAGGCACGAGCACAGCCAGGGCGCCCAGAAACAGGCACCGAGAGCGG | This study | |

| | | | |
|---|---|---|---|
| | | TGAGATCATCAGGAACAGAAAAAGATGTCATGCGATCCGGCAGCCGTCGACACCG<br>GGAGGCCTGTGCGTGTGATGCAATCGCCTGGTCGGATTCTAAACTACACCAGTCT<br>ATTCTCGTCTGGAAGGAAAAAAAAAGATACTTAAACCAAGCCTCCATCTCGTTTG<br>CCATCTCGCTCTTGCTTTCTTTCCCCCACCTTCTCTGCACACTCTTGTATCCACA<br>AAGCAGCTCATCACATATTCCATATCCATACCTCCAACCTATCGTTTTTTGTTTG<br>AACC | |
| $P_{HP10}$ | Promoter | TGATGGGGTTTGTCTCGCTGTTGTATATTTATCTTGTACTGTGAGTTTCTCTCTC<br>TTGCTTTTCATACCTCAGGTTGCGCGCTTATCAGATGTAGCACTTCATCAATAGA<br>TATCATTTAACCCTAAAAGGAGTTTATCACGTTTATACCCGTTCCTCTCTATTTT<br>ATCACCATTCGACCAAAGGACGATATTTCCTTCCTAAAGGACAGTTTGACGATTC<br>TTCCTTCTCCTCTTTCTTTTATGTTCGCATCTCGTTCGGCTTCATCTCGACATGT<br>CCATTCTCGTACAAGAAAATTCTAATATGACGCAGTAGCTGGTCGGCTTTATCGG<br>TTTCTTCCGCCAGTAAAGTCGGCGGTGTTCGGTAAAATCCATCTTCGGCCATATT<br>CGGGCAACCATATGGCATTTGTCGTTCACCGCCATATACTAACAGAGTAGAAACA<br>GACTGGAAGCTGATATCATGGTGACTCTATATGTTCGTGTGTCCTGTCCAGTTGA<br>CTTGAACTTGTTCAATCGAATCACATCATGAGGGAGGATAGATGAGGAAGAGTTG<br>GTGGGAAATGGCGAAGCAAAGAGAGATTGGCTGACGATGTCAGAATATTCAGCTC<br>TATTCTTGGGGATTCTCACTGATACCGTTCACTGGTGGCTCGTTCCATTCAGCCA<br>GTCATTTTACCGACTGACGAGCTTCTCACATCTCATCACTGGCCATGTGTTTGAC<br>TCGAAATAAAGCCGACAAAGGCGGGAACCCTTGGTTGAAATCATATATCTAGACG<br>CTATCCATCCTCTTTCTTCCTCTTCTCCTTCTTACTCATCTTCTCAATCACACAT<br>AGCCGTCTTTCTCACCAAGCCGCTCTAATATCAACACTTCACCCCAAGTTAAATC<br>TTACCTAGCCAAAC | This study |
| $P_{HP9}$ | Promoter | AGTGTAGTTTTAGAACACAAAAGGATCAAAGTTTCGTTTCTCGCCGTTCAGGGTT<br>AGAAACGTTTGGGTTTGTTTGATCCCCATCTTCAGTAAGAAAGAATGAACAAAAA<br>ATAGCCTCGGACCAAGTATTGTACAAGGAAGAAAGAAAACAAGAAATCATATGTC<br>GTTGTCATAGACACCATCGCAGTCACAGCCTTCACAGCACCACATCTGTAGGCTT<br>GTGGTCTATCGGATGTACAGCATCGCCTTTGACGCCTGTTACAACTCTGTCGTCG<br>TTGGGATTGCTGTGATTGCCTTTGTTCTCTTTGCTTGCCGTTCGTGACCTCGGAT<br>GGTTTGTTGACAAGCTTGCTCCCGAAAGAACCGGCTTTCGAATGTTCCATCGCCG<br>TCGGCCACAGCAGCGGCTTACACAGCAACACACACCTTAAATCGGACTCCTTATT<br>TATACATGTTCAATTTTATATTTAGCATTTCGTCAGCGTCCCTTCTTTCACATGC<br>GAGAATGGTATAAAATACCTCGACCGAACATGCTCCATATCGTTTCGATTCCTTC<br>CTTCCCCCATCTCATTCGCTCTTTCAGCTTACTACCCACATCTACTTCACTAACT<br>CCTCACCCAACCTTCCAACAGACACATAAATCACAATC | This study |
| $P_{MLRQ}$ | Promoter | AGACTTGGCATTCAAGAACTGGCAACGTCATCCATTGTCATGAGAACGTGTTGTA<br>TACCGATTTGACGAAGAATAGAGCATGGACTTTTAATAATGAACCATAGGGAGGA<br>AGGGCTTCGATGAATTCTTCCTCATGCGGTTAAAGAAACGGCTGTTCGCTTACCT<br>AATCGACTTTTCGCTGAACTTCCAACACCTTCCTGCAGACTCGTCGACTCTCGCG<br>CGTCATCCTCATTCATCCTCTGTCTTTCACGTCTTTCTCATCTGTTTCTTTCTGT<br>TCATCTCCGTTCGCCTACTGTGTAAAGGCTAACTAGAACATCGGGAGCGACTGTT<br>CATCCGCCATACTAATGCTTATCCGTTGAAGAAATGCTGACAATTTGTATAAAAC<br>ACCATGTTTCGCTAATATCTTTCCTGCCCCTCATCACATCATATCAAACTGACGC<br>ACACTCATATCTCCTCGCACTCACCTTCACATATTCTATCTCCTTCTCTATCCCC<br>ATCGAACAAT | This study |
| $P_{CYC1}$ | Promoter | AGTAGAGAGAGAGAAATCGAAATGGGATCAACGGCTCAGTTAAGTCCCTCAAGAC<br>TGAGGGGACTTGGCCTGGGACCGTCGGAAAAGACCGCAAAAGAAGTCTTGAGAAT<br>CTAGGAGTCACATAGACCTATCTGATTGAATCTCACATCCTTCCGTCGGCAATAC<br>GCAACGATCTCTAATCAAGTGAAGGGCTTTGGGATTATACATGCTTTTGTTATAG<br>ACAATCATGTCGATTTCGTGCGCAACGTATTATTTCTGAAGCTTTCTTGTATCCC<br>CTGCAAAGTACAGCATGCATTTCAACCTATTTACTCTAGCCCTTTCCCACTGCCT<br>CTGATGGATACTTTACGCGGCGTTATAAGACCACCAAGGATGAACACACCCGGAT<br>ACATGTATCTGTTCAATCGATCTCAGCGTATTGACTGGAGAGAGCGCATACAAAC<br>ACTAGTAGATCAATCCAAACGCCAGCTCTCTGGTGATCTTTGTCGTTTGCTATGA<br>TGAGCTCCCTCCGTCTTTGTCCGGAACTGGACGGAGAGAACGGACGGATCTGGTT<br>GAACCAATCGGAAAGTAATTATATATACATGTAACTACCACGAAGGAACAAACCA<br>CGATGATACAGATTACTACACAACAGGGAGAGATTGGCAAAGCCAGAAACATTCG<br>ATTAAAACCCCAATCCAACGCTGCCGCTGATGAGGCACTCGATGAGCCGCACTCT<br>GACCATCACAGACCGACGGACAAGACCCCACGGTGGAATGTTATGCAACAGCAAC<br>CGCCGAAACGGGCGAATCAGGGTGGATGTTTTAGGCGACCGTAACCTTGATTACG<br>TAAAATCGCGGAAGGCGGGAATCGTTTCGACGACTGCGTGGCGAATAAGAAGCTT<br>GCAATAAGAAGCTCTCCGGTTTTTTTTCCCTTCGACCACCACCAACGACTTTTC<br>TTTCTCTCACTTTCATTCTCTTTCTTTTCTCGTTACTCTTTCTACTCTTTAATCA<br>ATCTATCAACA | This study |

| | | | |
|---|---|---|---|
| $P_{E3}$ | Promoter | CCTGAAATCGAGACAGAAGAGCAGGCTCGAGACAAGAGTGGAACAAGACCTGGGGAAGATATATACAGGAGATTGTGAGAGACACTTTACGAGGATTACTACAGAAGCTCTATATCATTGCCACAGAAGCTCGATGTCGATATCTTGGATGCTGTACATCCATAGGTCGGGTACTATACGGGTACACCATCGGGACACCCACAGGCAGTACGGATGATACATTGAGAACACGCTTACGACTAGACTGTTTATATGTATATTGACACTCTATACACACATACTGTGACACGCACGGACGCCACTCAAATACACATCGATACTACTACACACACATTTCCTCCAAGTGCACTCAGTGTATGAACAGACCCAAGATCGGGAATGTGTTCCGCTGAGTCGATCTCGATCTGGACGAGAGGAGAGAGTCGATGGATCGGGAAAATGTAATTTTCAGGAACAGGCCATCCGAGGCGGCGGCGAGGAGGAGAAAAAAAAGAAAGGCAATAAGGCAATAAGGCAATAAGGCAATAAGGCAATAAGGCAATAAATCAATATGGCGGCGCCGGCAGAGAAGAAGCTAATAAGCCAATAAGCCACCGAAAAACACCAGGGCATCAGGGCACTAGGGCGCCAGTCATTCTAATCCACCGGCCGGTGACGTTGTGGATGATCGATCGAGCGAGCCAGGGCAAGCCTGCTGCGCTTTTAACCTCCAGACCTCCCACCGGTCGATCCCAGTTCCTGAGCGGGGAGAACGCAAGAGGTCCGTGCGAGGGTGCGTGCTCGCTAATTCCGTCTCTCCAAAGGAACACCACCACCAGGGAAAGCTGAAGAGTTTGTCGTTTCGTCTCTTGTCTTGACTGCTCTCTGTCGTCCATCATCATCATCATAGATACTCACACTTACATTCCCACTCACACATCATCTTACTCTGTACAACGATACCATTACAAGCAGGTTTGACTCTCTGCTTGAAAAGATTTCGTGAATTCTGCCATCCTCGC | This study |
| $P_{RPL6}$ | Promoter | GCTTGATGGAAAACGCTAGCAGCAAAGAAAGTTGATGGATTTGGAGGATCTCCGGTTAGTCACCGCTGCCCAATGGGCGTATGACTTTTATAGTTACATAAAGGTCAGCTTCCATGTGTTGGCAGCTCACGTCAGCCTGTGACCAGACTTGCTAAGGCTCAGGCCGCATAACCATCGATGGCGCGTGCCGCGTAACAAAGCTCGCCGGTCTGCCCGCCCGCCGTTCGCTCGGCTGGGTGAGAAATTTAGGGCTTTTCCACCTCGTCTTGTCAAGTTTCTCTTTGCACCTCCGACCCCTTGTATTTCTCGTCAAGGTTAGTTAACCCTTTAAAAGTGTTTGGAAATAGCTCCCCAAGCCCCTCAGTATCCATCAGTCCGCACATATCCAGCCATGCCAGTTCTGTATCTTCTTTGCAGGAATCATGAGGAGAGGGGAAATTTTCGTCGGCAGACTAGTTGTCCTAGTACGGTAGCAATGGGTTGAAAACCGAGCGAGGTAGTACGGGGAGAGGTAGTGAGGACATTGGGTGGGAAGCCCGATGGGCACCGACAGAATTACTTGGATTCAGCGAGTCAGACCGAGAAGATCGACATCGACAGAGGCAGGCCTCTCGACAGACGAACTGGAATGGATTCACGATATACGAAGAAAAATGACAGAACTCTGGCTGGATGGTTGTGCGGATCTGAGGGCTTTGTGAGCTGTCCTGTTCTTTTCATGGCACAGTACACTGACGGTTTCGGTTTTTCTATTCTTTCTATGCTTCTTCTCTGGCGTTCTATCTTTGTCACTGGGAACCTTCAACTCTGATGAACGATCTACGCTCGATTCTACTTGTTGTTCCCCATCCCTCCCATCGTTTCACGTCGGCCTCGCGACAAAACTTTCCATTTCTGTCCCTGCAATCGCGTTCATCTTTCCCGAATACCCACTTCGCAACCCCCAATCACTGCTTCATCTCCGTTCTCTCTCCTTGTTGCTATTGTGTTGGATTCAATAG | This study |
| $P_{CCP}$ | Promoter | AGTGTACAGAACATACAAACGCGTGTGCGTCGGGCTTGGATGAGTCTGTAGTCTTTTGATGAACGATGAAGGAGGTCTATGTCGTATTCAGATCTGTTGTTGTCCAAAGCTTCATGTGAGAAATCTTATAGGATCTGTCCGTGTTCGACGGGGTCGCTACTCAGACAAACCAGAGTGTCCTATTGTAAATGCTACGGAAATAATAACAAATACAGAGGGTTGATGGTCAAGAACATCGTCCATTGTGGGGGACACGACATTCCCGGTGTGGGGGGTTTTAAACTATTGTCGTCATATGTATGTAGAATAAGAGTAACCCTAACTTTCCCAATGTGGAAAAACTTGCAGTCAGATGATTGATCCTTCCGGTCATGCCGGAGTTCTCCAGCCTTAAAGGCAGCGGGGTGAGTAAGTAAATGTATCAACTCACTTCCTTCTTATCCATCCATCTAAACCTACCTTACCTTTTGTGTCCGCGAAAATAATCAGTAAT | This study |
| $P_{RP}$ | Promoter | AGATTATTCAGCCTAGTCGTCCGCTGGCTGCTCTAAGCTGTCTTTTAATCCTGCGTGGCTTCCCTAGTGACTCTGGTTCTCTCCGGGCTCAAACTCGGAGTCACAGGAGACTATGCACTCTTCTGAGAACGCTCCGAGAGCTCCCTGTATACTCGAACGCACAGTTTCAGCGAGACTATATAATATAATATAATATAAAACAGAACAAAAAGAGAATAGGTATCACTGAGGGCATCAGTGGCTTTAGAGCACTCAAGAGGCCGTGGCATTGGCCTGGTGGATGCAGAGAGATGTAAGCTGAGTGATCGTTTGGTGTAACCCTAACAAGGTACTTCATATATGCTACCTATATTCAAATAATATCCTCGATTCTCACGTGATCATACAAGAGAATGTCATCCTATCCTTTCCCCCTTCCATTCTCACTTTCGTTGCTTTCTTGCGCGCAAGGCATCAACTCTATCAAATACCTTTCGTTCGCTTCATTTATTAATTCGAATCACTCTTGAATGCTCTTGAATGACTTCCTCCTAGATCGAAATAGCCCTTTCATTGTCGCGTAGGACTCTTTAGCCTTAGGATCAGTAACACATGCCTATAATGATGATACCTCGATTGTTTGCGCATTGATCGAACGGAGCGCTGCGAATTTCTTGAGAGCCTGAAGATTCAATTCACTAGTTCAACCATCATACCTCCTCAGCGGACACATAGCAAACAAAACTATGTTATGGGTCTGAGAGAGAATCAACGTCGAAAAGATGTGTAAACAACTCTCGAGGCCATTTACCATGTTCAATACCACCCCTCTATTGAAAGTCTTACGTAATCATTATGTGTTACATCCGTGCAGCCTGGTTTTTCGTTACTTACACAATGGTCGGCTCCTGCAAGCATGAGTCCGTCAGAAAAATTATCTGTCGCCCCTCCCCCCCGCTCCCCCGTCTCTTTTTTCCTCACGTCAACAACAAGGACACCAACACTTGGTTAGTTTCGACGAC | This study |

| | | | |
|---|---|---|---|
| $P_{HP1}$ | Promoter | TCTATACCTCTTCTGATCACATCACTTGATAAGACAGAGTCTGCGCTCTACCGAT<br>TCTTTTATTTTCTTGATCCTCGATTTTTGTAAAGTCGAGTTGTGTCTGTCCTTGA<br>AGGACGGCGTTCTTTTTGTTTCTTTGAATACAGACAGCAGTGCTGCTTGTTTCTT<br>CCTCTTCTTTCTGATTCATTCTTTTCCTGTCTTACTCCGTTATGATATCACGTTA<br>TTCCCATCAGACTCGTCTCAAGTGAAAGAGCAGATATAAAGCGGGTCGTCCTTAT<br>TCTATACGGTATAAATGAAGCTTCACGTTCTTCCGGTCACTTGACTAGCAACATA<br>ATTAAAACCAACCAATTACACGTGGGATGGCGGGGTAAAGTTCATCGGCCCGGGT<br>TAGTCCCTGACCCAACAATATCAAGTTCCAGACCGTACTAAACGCCGACAGTGCT<br>TATTTCTGTCGGAGTCCTGGAGTCCCGGTGCCCCGGTGCCTGAGCTTGCCCGCCT<br>CCGCCTTCCTCGCAGGCTGAACATCGTTCGAACAAATGGCTCCTTCCAGTTATAT<br>CCTGAGTAAACAACAGAATGAATAGGACAAACATTCGGTACCAAAACACACATAT<br>ATGGCGCACACCCGTACCCTTAGCGCACGCGGGTGCACTCAAGCTCACACCCCAA<br>GACACGTTTCCTAAGGAAGAAGGCAAACTCAGAATACCAAATAAAAGAGGCCGAA<br>CACCTACTACCCGAAATCGTTAGGTTCGGTGTCGAAACGTGCACGGGATAGAAGA<br>GAATGTCCCTCCGATGAAACAGTGCGCGCCTTTCGGGGTCGCAATGAATGTCTGA<br>GACGGAAAACTCAACGAGAACGTTCAAAATGATCGGATGGGCGAGACAAATCTCT<br>CCTCTCTTTCCTTCTTTCTCTGTAAAATACACCCACTCTCTTTGCTTTCCCTCGA<br>TTTCTTCCCCCTCTTTCGTTCCTCATCGAATATCCTCCAGAAAGATACCGACCCT<br>CTCCTACTCAG | This study |
| $P_{ETFQO}$ | Promoter | TAGACATATACATTTTTTCTCTTTTCTTTTCACCTCATCTTGCTCTAAACAAATC<br>CTTCTTCTATAGACTAGCTATGTAGATGTCGAGCTGTATTTGTCCATTCAATCCT<br>TTCTATAAATGATCATATCCGAGTCATCTTTTTTTGGTGCTGTGGCTAACATGAG<br>GTCGACGGTCGATTGACCAATCAGAACACCGGTTTTGATCCAACCGCCGGGGCGC<br>CTCCATCAAGTTTTGGTCATTTTCAGTGTTTCGTTTTTCTTTATCTTCCTCTATT<br>TCTTTCTCACCCGCTTGCCCGATCATTCTATCCACCTTCACACAAACTCTGAAAA<br>GAACAAACACGGATCGGTGCATACACGGATTAAGTGTTATCTGATCCTGGTGAGC<br>ATCATTCATC | This study |
| $P_{HP2}$ | Promoter | CTTGCAAGAAGTCTTACTCCTTCTCATTTTCTCTCTTCGCCCTCTTCACAGCGTG<br>GCCTTGATGACAACCAAATATGGAAGGAACTGAGATTTTTTCCTTTTTTTTTGCT<br>CTCACTCAGCTCAACCAATATTTTCCTTGAAAGGACAAGTCGTTTTGACATGACT<br>ATCGTGTCATGTCTTTCTGCTCAGATGACGCGTCGCTGTGTATAGAGGTAAAGAA<br>CTTAGGGTTAATTCAAGATTTGGGTCTATGCATCTAATACTACATTGCTGTTCAA<br>CATCCTGTTGGCCGATATGCCAGGATCTTATGTACTATGTAGCAAAAGCCTCTGA<br>ATAGAAGGATAACAAACAAGCCAACTGGCTGTTGAAGGTCAGAGAGAAATCTCTC<br>ACAGGCCCAAGGAAACAGACTTACCCAAGTATAATTGACCAGGGGGAAACATGAA<br>CTTGAGATAGGGATCATTCCATGAGTAACTTTGAGGCCAATAATTTTACATCCGA<br>AGCTATATTTTGCATCCAGATGATCTGCGTATGTAACCATAAGCACATTGGTTAC<br>GTTTCATTTTTCCGATCGTTCCCCCGATATGAGTTTTGGCCATCGACTTTTTCCT<br>GGCTCAACGCCAACTTGGCTCTGTTCAGACCCGCCAGATCCCTTGATTCCCTGCG<br>TTTTATGGGAGAAAGTGCTGCTAACCCTGGGCAAAGGTCGGATCATTCACCGGCC<br>TCGTACTCTGCCTGTATGTCCTGGGGCTGTCCGTTAGGGAAGCTTTCCTGATACC<br>TTTCTCTTCTTTTCCATTTTGTTTGTCAACTGTCTTTCGTTCCCTTCTCTGCTCA<br>CAACCATTGCAAACACATATACGCCCATCAAGGAACAGCACTTCTTCCTCTTCTC<br>TGCTCAAGAATCCATTCTGCTTTGATTGACCACTACAGTTCTAGGCTGGCATTTC<br>CTCTATATTTCCACCAACGCCCAAGACTCTTCATAAGAGAACCTTGTCCTGACGT<br>TCGTTGTTCCG | This study |
| $P_{GYG}$ | Promoter | CAACGCACAAACCTAATACTACATAGAAAGACCGACAGACAGAACGTAAAGAAGA<br>GGAAGCCAAAGGCTGAACCAGTGGGTACCGTTCAGACAGCATTTGCGTCACAGAC<br>AACCTATACACGCTCAAGGATTCAAATAGAAGAGAGATACTATGATGTGCAATAC<br>CAAAGACGGAAATAGCAACGAGATGGATGAATCTACTCAGTGGCCAACGGCGACA<br>GCAGAGTACGCCCGAGTAAAAGAATTGCTTACGCCAGCCCCGGAATGCGCCATGC<br>TCATACCATGGTATACCATCATACCTATGCATTCATACCATACACTTTCCATGCT<br>GCCATCTGAATCAATTTGTCCGCTGCGCTCTCAGTGTGCGAAATTACTCATCTGA<br>TTTCGCTGCTTCCTTCCTCTGCCCAAGATGGACCGGCCGGCGTAATTGAGCTGTG<br>TCACATGCGATCAACATCAAAGTCAGCTCCAAGATTAATCCTAAAACAGTGCTGA<br>TGCAGACGGAGGCATATATGTCGTTGTTTTTGGCATGCTTGCGCACCTGAAGGTG<br>CACTACATCTTTCATGACGAGTGACAAACTCTCTTCCCCTTTGCTTCTTTCTTTT<br>CTCTTTTCGTGTACCATAAGCTGTTCTCTCTGAGTCAGGTTCTGCCTTGATCTGT<br>ACATATAGGTGGTTACATCGAACACGTCTCTTTCCTCGTATCATTTACCAACTAT<br>TGTCTATCTGTCCTTGTCCTCTTGCAGACCTAGCCGGGATCACCGAGATGGGATT<br>GAATGGACGCACCCCAACCGCGGCCCAAGTTGATCCTATAAAACCCCCAAATTAC<br>CCCATGGCATACCGCGATCTGGGTCCACACACGTGACGACATTGCTGTCACGTGA<br>CCTGCCGATCGATCGCCGGATATGACCGCTTCCCGAGTTATCGAGAGAAAACAAA<br>AGTGAAAAAAAAAAAGAAGAGAATCGTCCTTCAGCATACACAACCATCTCGATCA<br>ACCCATCTAACA | This study |

| | | | |
|---|---|---|---|
| $P_{SDR}$ | Promoter | ATCAGAAGGAATACAGTGCGTTCGACGTGGATGGAATGAATTGAAACGGAAAGGG<br>ATGGTCGTCAGGAGGAGGATGAGCAAGACGCGTTCACAAGACGCGTGTGCTGATA<br>AATTACATTCTTATCTAATCATATCAGCGTGTCTGTCGATGACGCTTCAACTCCC<br>GATCTTCGATCGGTTGCGGACGAACACGAACGCTCAATGGTTTAAAGAGGGGAGG<br>AAGAACAAGGATGCGCTCTCTTTTCTCCTTTCCTATCTCACCCACTCGCCACATC<br>ATCTGAATCAACACATC | This study |
| $P_{NUCR}$ | Promoter | AAAACACTGAACTTTCTGCACCTTGGTCGGGCGCAATAAATACAATAAGCTGTTG<br>AGTCCAGAACGAAGGCGGTTGTGCAGGTATATGGAATGAATGAAGTGTTGAAGAT<br>TGTGGATGATCCACAGAAAAAGACGTGGTATGCCGAGGTATGTCTCTTACAAGAA<br>ACCTATCTATATCTCTCAAACCAATAGAATTTTTTTAAACCTCAATTCTCATTCG<br>GATACACTGTTAAAGAGTTCGTACCGGCTTGCTCCTCAAAGCCACCCGTTCTTCG<br>AGTTGACCCGGAAAGCTGCCAACAGGATTGGTAAAGTCAACCGTTCTGGTGAGTT<br>CCTACTTCGGCCTTCAACTGATGAAATCAGTCTAGTTTACAATCTATCCGAATTG<br>GCATCTCACTCTTTGATCTCTCATGTATCTTTTCTCTTCTTAAACCAGATGAGAG<br>CGAGCCGGTGATGGTGCTTCATAGGGCATTGGGAGAGATTCGGTCGTTTGTGGGT<br>GAGAGACTTCACCCATCTCAGTCACGTGATGGCTACACGTAACAATGTCATGGTT<br>ACAACTTTCAAAGACATGATTTTGATTGGTGCAAGAAAAAACAACGATTTTTGAA<br>AGTTGCCGttctacttcttctttaccttctctctttcccttccttctgttcgtct<br>ccttgtctttcGTTACTCTACTGCTTGTTGC | This study |
| $P_{HP3}$ | Promoter | TAGTTAGTTACCTCCTACAAGGTCTGATACATCGGCTTTGCTCTACCCTTTCTTG<br>TGGTTTGCGTTTGTTTATCTTCTGAATTGGTAGCCTTATCTTTTGATTTTCCATC<br>TGCATCAATTAAAGGCAAATCAACTGGGGTTTACTCCCATCGTTTGTCTCTCATT<br>CTATCTTATTCttattttgtcttggcttgttatgttttgttttgttttgttttgt<br>tttgttttgttttgttttcttttcttttcttttgttttctttCCTTATCCCTCGT<br>CCTGTTGTAAACGTTTCTTTCCTCCTGGTGGTATATATGATTTATCCTCCTCCTC<br>CTCTTGATTGTTTTCTTTCTATCCAATCGGGCCGACAGGCAATGTTACAAACAAA<br>ACAAAAAACACATGGGGTTTAAAAGACCGATGAGACTGTTTCGAACGGAGATTGC<br>TTGTGTGTTTCTGTGGCTGAGACGGCTGAAAGCTGTGTATGTGTATCGATTGACG<br>TTGGCCAATCGATTTTGCACGTGATCTCGAAACTGCGCGGCGGCCCTCGTCATCA<br>CGATCTCAACAACCGTCTCTTCTACTTCCATCGAACCAGAAGAGTGTGAACCATC<br>CCGTTTTTTCTTCGCCAGTTCTATAAACATGGGTCTTATGCTGCCCGTCTCTTTC<br>TTCCTTGGATGCATTCGCTTTCCATACATAACGTCCGTTCAAGATAGACTACCAA<br>AGGTGCTTCTTCAGTATACCTAGCCCAGGATCTCTCCTGACTCTGCTTCGCTCGC<br>TGGTATATGGAAC | This study |
| $P_{RPL19}$ | Promoter | GTCTCAGTAGAGAACGAGGGGTGAAAAAGGAAGGAGGAAAAAATGATCACTTCGC<br>CCTTGGTCGATTGGTCGTATGGAGAAAATTTAAATGCAATCGGACGACCCTTCAC<br>TCGATCGATGGAGCTCATACCATCGAGAGGGTCAGGTCGTCAGCGTTGAGGGAGG<br>ATGAGAGGAAAGGAGGAAGAAAAAAAAACCCCAAAGTCTTGCTCTCTCTTTTACC<br>TCGTCCGTTCCACGAACGAAACAGAGTGGCAATAGAGAGGGCAGTTTTGGACTCT<br>TTCTGTGCATGATTGGAACGGAGCTGTTCAAACATGATGGCAACGAAACCCACAT<br>CGGTATCACACATTAACGCCCCTCGCAACGACCTCACGGACACTCATACACACAC<br>ACCCAATACAAACATTTTTGTCAGCTTGATTTATTCCCAGAGCTTCCTACTGACA<br>TGCTTGCCTTTGTCTATTCTTCAG | This study |
| $P_{Con-6}$ | Promoter | ATGTACAGGACTAGATGGGTGATTCTACACGGATAAATGTACTGATGCAATATCC<br>AGACTCTTGAACGTGGAGCGTCGGAGAGAAGCGATCAACTAATCGTAGCGAGAAA<br>CACAATTTGATCACATAGAATATAGACGCGCCTGCTTTTTCCCACTTCGCCATCT<br>CTTCACTCCCGATCTCGATGAACTAACAGATGAAAACCATTCATCGAGTAATCTC<br>TGGCCATCTACTCTTCCTTTTCAATGTGACCGATCGATCCTGCCTGATCGTCTTT<br>TGTTTGATATCGTGTGACCTAACTTACATCACCTCTTTTGTTATCGATCATCCTC<br>ACTGACAGATTAGACCAACCCCGATCACACTAGCAGTTCTCATGCATGCAGATGA<br>TAATCATTCCACTCCATCATATCACGTCACATCGAACACGACTTCCGATCTCGAG<br>AGCCCTGTTTGTTATTCATGATGTTGACGACAGGTCTGATCCTTCGCTTCAAGGT<br>CTTGTCAAAAGTCTATATAAACGATGTGTGGATGGTCTGGTCTTGTGCACTTATC<br>ACCATCACATATCCTCTCTAACTGAATCCAAATCGATCCAACCAAACTCTTATAC<br>GTTTCAACCTTACACACCAAC | This study |
| $P_{SAM}$ | Promoter | TTCAAAGGGTAAAATATCACGTCGTAGGAAAGCCTAACGAACATCACAGACAGCC<br>GTCACGGTTGTATCTTGGTTCATACCTCACACCTTCTGGTGATCCATCAGCCAAG<br>CCCCAAACTTTTCTTTCCGTGCGGACATGGATATAGAAAGATCAACTAAAATTCA<br>TATGTTCTTTCAACCCATACACCAAATCGAGATACGACGTCAAAATGTGTCGTGA<br>TAATAACAATAAAAAGATGAAAGCTAAAAAACCCAAGTTAAACTCTGTCCCAAGA<br>TCTGGAGCTTGCAACGAAAGGAAGCTGGTCCAGTATCATCGTCGAGGAGATCATG<br>ACAGTATTGGCCTCACTGATTGGGAGAGTACAGTCTATGATAGCTAGTTCGCAGG<br>GATCATGTTTGTTTATGAATCACAATATGCGAATTTAGCGCAAAGTCCTATAAGT<br>GGTAATGGCAAGAATGTTTCGAAGAAGGAGTATATGTCGACAAGACAAAGAGAAG<br>ATAGGAAAGGACTCAAGACGAGGAAGGCTCTGATCTTGATATCCAAGAATGACAT | This study |

| | | | |
|---|---|---|---|
| | | GAAATGATAGGGTCGATAAGAAGATAGAGGGAACCCGTCCGGTAGGTACGATCAG<br>AAGAACTATCTGGGAGGTTCTGGAATGAGAGGATGGTGATGCAGTGTATCTAATA<br>CAAGCATCAAATAACGACAGTATCTGATAAGTTTACCCATCCCACAGGCGTCGCC<br>CCGGTTGACGCAGATCAGTCAGAGGGAGAGACACTGCATATCGGAGGGACCACAG<br>GAGACACGGGACAGAGCCGCGCGAGATCATGAAATAGAGGGACGATCAGCGATTG<br>TCTCGGCTGAGGGTTACTATCACCAAGCCTAAGGCTTATCATAGATCGGTGCACG<br>CGACGGTGAAATTCCGATGGAAATCCCATTAATCGGTCATCCAGATCGAGTATGT<br>ATTCCCATCAGATCACTAGTTTCCTCTGAGTAATCGTACAGTGTCCTGGTTCTCT<br>TGCTCTTCATC | |
| $P_{Nth}$ | Promoter | ATGTTCTATTGTTCTCTGTCATTACCTGTGGCTAGCTATATGTACTACCCCGAGA<br>ACTACTCTTCTTGATTTCATCTATGATCGTACTCCCTGTTGTGCCTCGATGTATA<br>TTATGAGAGAATTTCTGGCTGTTAATATACGTTGCTTGATATCTGGAACGATTCT<br>CGTTGATCTGGTGTCGTTCTCCTCCACGTATCTAATATGGTCGACATCAAAGATC<br>TTGTGAGCGAACTGTATTTTTAACATGAGTGTTGTTCATGTTAATTTCTTTGATA<br>GTTAGCCCAATAAATCCTCTCCTTCTCTCGTCATCATCAAGAGTCAGATAGAACT<br>GAAAGTGTCGAAACCCTATTCAAAACTCGCGATAAGAAACTCTTCTGAACATAGA<br>TCAAGAAGGAATATCTCAGATGGTCAAGCCTAACCAAACCTATCAATGAATTCAA<br>GAGTTTGCTACATCTTATCTTATCTCTATCACGGCGATCGGGAAAATGAACAATG<br>ACGCAAATCTAGGCTTAAAATCTCGATCCCTTGAGCTCGTCCGGGTACCCCTCTT<br>CTTACTTTGTACATCC | This study |
| $P_{TR}$ | Promoter | TATTCTTCCTCCCTAATGGAATGGATGGTGAAGGCAAGCAAAGACGGAGCAAGGA<br>AGAAAGAAGAAAGAGGGGAAGAAAGATAGATCCAGAGAGATGCCTCTGTTCAAAT<br>AGACTAAACTTCGTCCTGCAGTTTATCTGATACTCTGTTTCAACCGGCCCCAAAT<br>TTGAATCTGATTGAGAAAAATCGTGAAAGATGAATGAGAAATGCTCAGACTTCTT<br>CTTCGGTTAGCCAATCAGATGAGCCTGACGGGGTTTTTACAATCCCCAAATACTC<br>CCGAGCGACTTCGCCGGGGTTATAAACTCTATTCACATCTGAGCTCGAAGGCGaa<br>agaagagaaataagatggtgatgagtagaaaatgaagaaaggaaaggatagggaa<br>aaaagaagagaatgggaaataaaaattaaaaaaaaagaagagaaGGAGAAGCGAT<br>CCGACCCATGCTGAAGCTTGTTTAGTCAAGACTGAGCGCCAGAACAAAGCTCAAC<br>GAATTCTCATCCTATAGAAATTCACGACAGAAAGACGGCGGAACGACTGTAGTAG<br>GGAGGCATGTCTATGTGAGTTCTATCTGATAGTCAGATCACTTGCTCTATCTGAT<br>TAACTCAAGAAAAGGAGAAGAGTGAACGAAAGAAGCAGGAGAGGAACAGGAATGT<br>CAACTGGGATGATAACTTAATCAGATTACATCGCTATCAGTCAGTGGCTTTGTCA<br>GTTCGTTGCAGGTTCAAGAACCATATGTTCTATAACGGTCAGGAGACATTCTTGA<br>AGGCAGCCAGCATATACAGTCAAGTAGAAAGAAACGAACAAGTCTCTTGTCATTG<br>ATCTGAGGATGTCTCCAACCCAAGGGATGACAACGAGGCAATCAGAAATCTGGGA<br>AACACCAGATCGAATATGACTCGGGTGAGGATTGGGAATAAAGAAAAAATGCCAG<br>TAAAGTTCGAGACACATAGCTCAGCTTGAGACGCAGAAACCACGACAGCATCATT<br>CTTGACCCAGC | This study |
| $P_{HAP3}$ | Promoter | TATATTTATGCAAACACCAAGCACACCAGAATAGATTTCAAGACAAACTCAACAT<br>CTGTAGCAACGAGAGTCAACGATGTAACATGAAATATCTTAGTTGGACTGGATGG<br>TATGTAGGACTGATAGTATGAGATTGGATGCTCTTCACATTGAGAAGAATTTGGC<br>TTCACCTGGATATAAAGATGCTCTGACACACTTCTAGAACACAGACGATCGAAAT<br>ATGGTTAGATATATAACAGCATATCGTCAGTATATCGACTGGTATCTCATCGCAA<br>TAGAGGGATCTGGTATCAATCAGCATAAAGTTAGATCAAAGTGTGTGCTTATGAA<br>GCGAGGAATATTGGTATATTCAGCGATAGTATCAGCCGGGCAGGTATCGTCAGCA<br>GAGCAGGTCGAACAACACCATTGGCGTGTATATGACTATAGAATTACATTCAGCA<br>CTATGTTCTGTTCTGCTATGGTTCTCTCAAATGAGGATGAAAACTCTCCGAGTGC<br>AAAAGCGGACTCTGGATGTTCTTACGTCGTGGGTCACATAATCTGTCGACGCCGT<br>CTATCCGACGGCTCTCCTTACGGCGTAGCCTGACCTATATACACATGTTGTAGAT<br>CCTTCTGGTAACATACTCTATATCCATGTCATGACCAAACATGGACGGATGAGCA<br>CTTAAAGTACTCAAACCTGTCCGACCTGATCTGACGGCCGAACCCCAGAGGGTTT<br>AGAATCAAATTCGAATCTGGCTGACTACCATTGGCCAGACGGCGAAAACCGATTC<br>TCGAACGTTTTGATTTGTCCCCCTCTCTTTTTTCCAAGGAAGTGGTTTCTCTTCT<br>TCTTCTCATTTTCTATTTCTTTGCCGTTTGTCTCTTTTCGCTATCTTTTCTGTCC<br>GACCACAGCCTCCTGGATCTCTCCCTCTTCCAGATCAATGTCATCACCGTCAGCA<br>GTGTCATCCCTTTCCCCAGAGACAGACTCAGATGTCGAGCCGATAGTTCCGGTCG<br>TCGGGAAGCAA | This study |
| $P_{MSS4}$ | Promoter | ATAACCTTCATACACTCCACAATCCGAAATGTCTTGCCTGTTTTGTTCTTCCTCC<br>GATCGGCTGAAGGGATAGTCTTTGAAAGGTGATGTGGTCGGTGACGACCGGCCTA<br>TTTTTCTTTCTTCTATTGATTGTTCCAAAGCGTACATATTTGTAGCCCTGtctcc<br>tcttctctcctctcctctcctTATCGATCTGTTTGTCCGTTTATATAGCCCCTAG<br>GCTTGAATATTGTAATACGTTCATCTGTCTTTTCATCTAGTCTATCCAATTATTA<br>TACATTGCAGGGTACCCACAAAATCACGTGTTGCTTGCCAGCCACATATTTGCAT<br>CCACAAGCACAGCCCACTTTCGTTACCCCGGCCCGTATAGAAACGTCATAGTAGC<br>CAATCAATTGTATATTGGATATCTTCATCGTGTGTTCCATCATCGATCGTCGATC | This study |

| | | | | |
|---|---|---|---|---|
| | | GGGCCCGTTCTAGCCTGCTTTCTTCATCCCTGGCTTTGTCTTGCCGCCAGAGTTC<br>ATGCCAAAGATGACGATGGAGC | | |
| *natR* | Gene | ATGGGTACCACTCTTGACGACACGGCTTACCGGTACCGCACCAGTGTCCCGGGGG<br>ACGCCGAGGCCATCGAGGCACTGGATGGGTCCTTCACCACCGACACCGTATTCCG<br>CGTCACCGCCACCGGGGACGGCTTCACCCTGCGGGAGGTGCCGGTGGACCCGCCC<br>CTGACCAAGGTGTTCCCCGACGACGAATCGGACGACGAATCGGACGACGGGGAGG<br>ACGGCGACCCGGATTCCCGGACGTTCGTCGCGTACGGGGACGACGGCGACCTGGC<br>GGGCTTCGTGGTCGTCTCGTACTCCGGCTGGAACCGCCGGCTGACCGTCGAGGAC<br>ATCGAGGTCGCCCCGGAGCACCGGGGGCACGGGGTCGGGCGCGCGTTGATGGGGC<br>TCGCGACGGAGTTCGCCCGCGAGCGGGGCGCCGGGCACCTCTGGCTGGAGGTCAC<br>CAACGTCAACGCACCGGCGATCCACGCGTACCGGCGGATGGGGTTCACCCTCTGC<br>GGCCTGGACACCGCCCTGTACGACGGCACCGCCTCGGACGGCGAGCAGGCGCTCT<br>ACATGAGCATGCCCTGCCCCTAA | Young, 2018 | [1] |
| *NLuc* | Gene | ATGGTTTTCACCCTCGAAGATTTTGTGGGGGACTGGAGACAGACCGCTGGATACA<br>ACCTCGATCAGGTTCTTGAACAGGGAGGCGTCTCCAGTCTTTTCCAGAATCTTGG<br>CGTGTCTGTCACCCCCATTCAACGAATCGTCCTCTCGGGAGAGAACGGGCTCAAG<br>ATCGATATCCACGTCATCATTCCTTACGAGGGATTGAGTGGAGATCAGATGGGAC<br>AAATTGAAAAGATTTTCAAGGTCGTTTATCCTGTTGACGACCACCACTTTAAAGT<br>CATCCTGCATTATGGCACTCTCGTTATTGACGGAGTTACACCGAACATGATCGAT<br>TATTTTGGTAGGCCTTATGAGGGTATTGCTGTATTCGACGGAAAGAAAATTACCG<br>TCACTGGTACTCTCTGGAATGGTAACAAAATTATCGACGAGCGACTTATTAACCC<br>GGACGGTTCATTGCTGTTTCGGGTCACTATCAATGGAGTCACTGGTTGGCGGCTC<br>TGTGAGAGGATCCTCGCCTAA | England, 2016 | [2] |
| $P_{act}$ | Promoter | GAGGACGATGTGTGCATTGATGATCCTGAGCATAAGTTAGGCGTACAAGAGCATT<br>ACAGCCTATATGTGAATGGAAGTGAGAGGTGGAAGTCAGCCTCCAGACGAGTTAT<br>GCAGTCTAATCAGGGAAGAGCTGGTCCAGAGCCAAACAAACTTACTTCCGGTGCC<br>ATATTAACTTGGATAAAGGTATAGGATAGTATCGATTGAAATGGCTGTGTCTGTG<br>AGAAGATATGACAGAGAGCACAAGATGGGATGGAAGATGCCTCTGTGGACTTGAC<br>TGATGGATTAGCTGGAGGTCTGATCGCACGGCTTAGTCAGCTAGCTTCCATGTGG<br>TTACGACTGCGATCGAATTATATTCTAAATTACAACCGTAATTTAGCCGTCAACA<br>CAACCGTCCGTGCCCAGGCTCCATCTCTTTATTTCTCTTCTCTTTCTCTTCTTCC<br>CTCCTTCTTCCCGACCACCACTCACTCTCACTCTCTCTTTCTCTTAGAACAAAAC<br>TCCCGTACCTCCTCCACCTTTAAAAAGAAACAGCATAGCCCACC | Hara, 2014 | [3] |
| $P_{adh4}$ | Promoter | GGTAAAGAAAAGAAAGTCGACCGTGTAAGAACGGAGGATCCAAGAGAAGGCGACA<br>TTCTGAAGCAACCAGACGAGGAGATAGATGACTATGGATTCACAGAGTGTTAGGG<br>TTAACGAACTCGATTAGTCAGCTTGATAACGCGCAGTAGCCTTCACGCGCAGCAG<br>CCACGTCATTTCGGCCCGGCGCAAACGGATGCCGAGTGTTTTTTTCGGGCCGTTC<br>GGAAGAGAGAGGAACAACGACGATGGCAGAGAAAACATCTTGGGATGGACAACTC<br>TTGCTGTTCACGTCAGAGGCTGAGGACCAAGGGCTGGGTATTGGGGTTTGAGATT<br>GACTATAGAGTGATAGTAATATCTGGAGATCAGAGCTGAGACTGATAAGACTGAT<br>GCTAACGAGCTGACGCTGATAGATTACCGGTGCTGCTGAAACTGCTATCCGGACT<br>AACGAGGTGGAAGATGACGGAACTCTTTCTTCGCCGGCTGTTCCATCTTCTTCTT<br>CGTCTTTCTCTCATTGCCTCGTTTTTCTTCCCATCGAGGGGCTCCTCAGAAACGT<br>CCGGTTTCCGACATCTTCTGTCAATCTTTGTTCCAGTCTGTGGTAGGTGCTTGTG<br>TGCTGATCCTTACGGCCTCTGTTCTTCCAGGGGTTTCCCGTGGGAAATCTGTCCG<br>GGAAGGGATCCGATTGATTCCTGACAATGTCCTTTCTTTCGAGCGATAGGCAGAG<br>GACTGGTTTATTTAATGTTTCCCTTTTCCGTCTGCTCGCTCACGGAGGATCCAAC<br>GTTCTTTCTCTTCCTCCACTTCCGTATCGTCCGGGGCAGATCGTCATCACAGAAC<br>GGAGACTGAAATGAACCATATAAAGCAGGTGGTGTCTCGATCTCGATCTCACTCT<br>CCTTTCTTTCCCTCACCATACAAACATCCCATCCTATCAAGCCTCAGCATCACTC<br>CTAGCTTACTCTCACAGAGATCCTATTCAACCAACCAACCAATAACACATCATCT<br>GAACCCCACA | Hara, 2014 | [3] |
| $P_{gdh}$ | Promoter | ACACGTGATATATGGCTGTACGAAAAAGTCCGAGAACCTGAAAGAACATCCCAGA<br>ACGTCGGAGGATGCCGATGCCGATGCCGCGCACGGACATTGGGAAACACGAGCGG<br>AGCGACCGAGGGGCGACCAGGGGGGGGGGACGGAGGAGTGTGTGAGGAGGAGTAG<br>GACTGAGGAGTGTGTGTGCGAACGGGGCCTGGTGTGTAAGCCCCAGGGACGGCTC<br>AGGCGCTTCCCAGGCACTTTCAGGGGCACCCAGGCACGGAGATATGAAATCAACG<br>ATCGACCCAACTGATAAGGAAATTTGCAAATCGCCCTGATCTCGTTCCTGACTGC<br>GCAGTTTTTCTTTCCCGTCCCTTTTTTCCCATATTTTATGTTTGCCCTGCCATCT<br>TTTTTCCTTTTCCTCCCTCTTTCTCTTTCTCCACCATCTTCATCCTTCCCATCCC<br>ATCTCATCTATCTCCCATCTCACCTTCTCCTCCACATCTTCCATCAACCATG | Hara, 2014 | [3] |
| $T_{act}$ | Terminator | ATCAACAAAGTCTTTCTATCCTTTAAGGCAGAACGGTTTCTTTTCTTACGATAGA<br>GGCACCCTTGGCGGTCTTTCGCCAGCGGTGGTTTTCTCACTCTTTTTCCAATATT<br>TTCACCGACGTTTTTGCTCGCGTGTATGTTTTTCTCTACTTGATGCTATCAATAT | Hara, 2014 | [3] |

| | | | | |
|---|---|---|---|---|
| | | CATGGCTTTTACGTTCCGTTGTATTATCATCTTCTGCGCCCTTCTGACTGAGATA<br>TCGAAATCGTTACAACACAAGACATATAAAAATGAGATTTATTCAACTACTAAAT<br>AACGATACATTAGACGTAATAAATAGATTCATAGACAGGCCCGAATGTTGATCGA<br>TCAAAGACTTTAAACGTAATTGCGACATTCTCCAAAACCCCAAGCCGAACTCTTT<br>CCATCACCTATTTGTTCCCAAATTTCGAAACATCCACTCTCTTTCTGAAGCTCAG<br>TATCACCTCGAGAACGGATCTTCAATCTTTAGGGCGTGTCTTTTTCCGGTCTTAT<br>CTTGAATTTGTTATCGTCATTCGATATAGATCTCCCTTATTGTATTCTATCCTTC<br>GAACGCGTTTTATACGTATCAACCCTCGACCATGCCGTCCGATTCATCCCATCGA<br>GCCTCACGATCTGGTTCATCAGACAAGAACAGGTCGAATACAGACGGTCCATCAG<br>CTCAGTCTTCGTCTCTGTTCTCCGGATCAGCTTCAGCTCCGGCAGGT | | |
| $T_{gdh}$ | Terminator | ATTACCCCGCCTCTCTCACATTCTTCCCCCTCCCCTTCCGCCCTTCTCTGCCTTC<br>TTTGATCATCTCAATCCTACTGTAATCTTTTTCCTTTAAAATTCGAGATTTGTAT<br>CCCCATCGCTCGTGTGGTTTGCCCTGCTATCTTTTCCTCTCCCCCCTTATATTCA<br>GTGCTCTCCCTCTTGGCCTCTTTTGTTTCTCATATGTCTTGGGTTCTCCTTCCTC<br>TGCTGGCTTATCTTATATCGTCTACGACATGATGATCTTCTTTCGCCATTTTCTT<br>TCCCCCTTGAACAACGGCTCTTGCGCGAGCTTGCCCACATGCAAATGCAAACATA<br>TATCATTTGATCATGATCATAAACATAAAGAGAAACACGATACAGCAAGCAGGGG<br>CATGGGCATACACAACAAGCGTTATCAACCAGAAAGAAACAAGTATATCCTATCC<br>GCGCTCT | This study | |
| 5’ int crtYB | Genome fragment for targeted integration | CAGAAGATGGGTCCACCGATAGTGGTTTGAAAGGAGATGGGTGGAATAGAGACAA<br>ATGGGGAATTCCAGTTTTGCCTTTGACGAGAAAGGACACTGGGTTGGAAAGAGAA<br>GATGGTACGTTCTTCTCCACCTTGAATGTGTTGCTTACTAGACATGTTTGACACG<br>CTAATGCATTTCTTTCCACTTTGACTTTTGAACTATGGTGGTTGGGCGATCCCCA<br>AAATCATTAGCTTCTACTTCAGCTCATTACCTCGATCTCATCTTACTACCAGGTG<br>TTGCATTCTCACCTACGGCCTCTTCTTTGTTCTCTCGACTGGGCCATGGAAAAGG<br>ATATTACGATAAATACATCACTCAGTATCGGTCGATCTGTGCAGGCAAGAATCGA<br>CCCGGTCCGAAGCTGAGTACGCGTATTCTCTTTTCTCGATACCCAACGGACGCTA<br>TTTTGTGACAGAAGGATGAGACTATCCAACAGCTCAAACAAACTAACGCTCTTGA<br>TTAATCACCCGCTCAACTTATTGCTCAACTCAGTTGGACTGGCGCTGAAAGAACA<br>GTTCTTAGACAAAAACATGGTCCCTATAGGAGAATGGGATGCGAATCTGGATGAA<br>GTGTTGGTTGGAGATCACGTGAGGACATTATCCGAGGACAATTAACTACTTAAGA<br>TATATACATGATTTATGTCGATCGGCATCCAGCCGGGGATTGATCGGCTGATGGC<br>CGGAAATGTGATGATGGTCGAAACTCGATCTCTCTTTTTTTGTTCATCTTCTCAT<br>CCCTCTTCTCTCTTTCTACTGACATCCATCTCCAACTGTCTAGATCAGTTCGGAA<br>ACAAGAAGTGGACACAGAGAGATCTTTGCTGAAGAGTTGTATTCCAGAAAGGGAA<br>AACAAAGGAAAGAAGCGCCGAAGCACATCACCAACTTCAGCAAGCCGGTCCAGCC<br>CGATCTCGGATAGACATCATCTTACCCAACTCGTATCATCCCCAACAGATAGAGT<br>TTTTGTCGCA | This study | |
| 3’ int crtYB | Genome fragment for targeted integration | GACAGCGGAAGAATACCGACAGACAATGATGAGTGAGAATAAAATCATCCTCAAT<br>CTTCTTTCTCTAGGTGCTCTTTTTTGTTTTCTATTATGACCAACTCTAAAGGAAC<br>TGGCCTTGCAGATATTTCTCTTCCCCCCATCTTCCTCCTTTCCATCGTTTGTTCT<br>TTCCATTTTTGTCGGTTTACTATGTCAATTCTTTTTCTTGCTTTTTCTTATCAAT<br>CTAGACAATTCTATAGATGTTTAGAATTTATACATTGACAGGTTATAGACCATAA<br>AGACTATATGGCTTTGGCTACTAGAGAGCAACCGCAACCGATGGTGGTCGATACG<br>CGGCCCTTTTGGACTTGGGTGCCCTCAGACCACAGACCACCAAGATCCCCGTGGG<br>TCACGGGCTTAGCTTGTGCCAGCCCCAGCAGTCATGACAGTCCCAGCCCCAGCCC<br>CAGCCTCACTAGCAACAGCGTCAAAGCCCGGCGTCCTTGCCAACTGGTTCTCCTC<br>CTGATGCAGGGCTAGGTGCAGCATGTTGCGCGATACGTCGACATACCTGTTCCAC<br>CAAGTCTGAGAGAACACAGTAAGCAAACATAGAGAGTAGTGAGCAACCGACACAT<br>CATCTTGAGCCATACAACACATGTCACTTACTCGACAGACGCTTCGACACTCCTA<br>GATCGCCCACGCTTTGCCTCTTCCAGAGAACTCGCCATGACACTTCGAGCACACT<br>CTCGACGTCCGATACTCCCGGACTTTGCGAGCCTGGATCAGTAGGTAGATCCTCA<br>GCTTGGCAAACAGCACCTTGACAGGACCAGGAGCCTGCCCACGAGAAGGCGCCGT<br>CGCCGAAGGCCAGGAAGGTGCGTGGACGTCCTGCTGACAGACGGACGGCCACCTT<br>GCGGTACGTTTGCTCGCGGCTGATGTGCGCGCCCCAGTGGAGACGTCTCCACCGC<br>CGAGCGGCGAAGAATGCCGCGAGCCAAGAGAAGGACTGATAGCGCGAGACGAGAG<br>CAGCATGGAG | This study | |
| ColE1 | Cloning replicon | GGCCGCGTTGCTGGCGTTTTTCCATAGGCTCCGCCCCCCTGACGAGCATCACAAA<br>AATCGACGCTCAAGTCAGAGGTGGCGAAACCCGACAGGACTATAAAGATACCAGG<br>CGTTTCCCCCTGGAAGCTCCCTCGTGCGCTCTCCTGTTCCGACCCTGCCGCTTAC<br>CGGATACCTGTCCGCCTTTCTCCCTTCGGGAAGCGTGGCGCTTTCTCATAGCTCA<br>CGCTGTAGGTATCTCAGTTCGGTGTAGGTCGTTCGCTCCAAGCTGGGCTGTGTGC<br>ACGAACCCCCCGTTCAGCCCGACCGCTGCGCCTTATCCGGTAACTATCGTCTTGA<br>GCCCAACCCGGTAAGACACGACTTATCGCCACTGGCAGCAGCCACTGGTAACAGG<br>ATTAGCAGAGCGAGGTATGTAGGCGGTGCTACAGAGTTCTTGAAGTGGTGGCCTA<br>ACTACGGCTACACTAGAAGAACAGTATTTGGTATCTGCGCTCTGCTGAAGCCAGT | Young, 2018 | [1] |

| | | | | |
|---|---|---|---|---|
| | | TACCTTCGGAAAAAGAGTTGGTAGCTCTTGATCCGGCAAACAAACCACCGCTGGT<br>AGCGGTGGTTTTTTTGTTTGCAAGCAGCAGATTACGCGCAGAAAAAAAGGATCTC<br>AAGAAGATCCTTTGATCTTTTCTACGGGGTCTGACGCTCAGTGGAACGAAAACTC<br>ACGTTAAGGGATTTTGGTCATGA | | |
| *ccdB* | Cloning selection cassette | ATGCAGTTTAAGGTTTACACCTATAAAAGAGAGAGCCGTTATCGTCTGTTTGTGG<br>ATGTACAGAGTGATATTATTGACACGCCCGGGCGACGGATGGTGATCCCCCTGGC<br>CAGTGCACGTCTGCTGTCAGATAAAGTCTCCCGTGAACTTTACCCGGTGGTGCAT<br>ATCGGGGATGAAAGCTGGCGCATGATGACCACCGATATGGCCAGTGTGCCGGTCT<br>CCGTTATCGGGGAAGAAGTGGCTGATCTCAGCCACCGCGAAAATGACATCAAAAA<br>CGCCATTAACCTGATGTTCTGGGGAATATAA | Bermand, 2018 | [4] |
| *kanR* | Cloning antibiotic cassette | ATGGGTAAGGAAAAGACACACGTTTCGAGGCCGCGATTAAATTCCAACATGGATG<br>CTGATTTATATGGGTATAAATGGGCTCGCGATAATGTCGGGCAATCAGGTGCGAC<br>AATCTATCGATTGTATGGGAAGCCCGATGCGCCAGAGTTGTTTCTGAAACATGGC<br>AAAGGTAGCGTTGCCAATGATGTTACAGATGAGATGGTCAGACTAAACTGGCTGA<br>CGGAATTTATGCCTCTTCCGACCATCAAGCATTTTATCCGTACTCCTGATGATGC<br>ATGGTTACTCACCACTGCGATCCCCGGCAAAACAGCATTCCAGGTATTAGAAGAA<br>TATCCTGATTCAGGTGAAAATATTGTTGATGCGCTGGCAGTGTTCCTGCGCCGGT<br>TGCATTCGATTCCTGTTTGTAATTGTCCTTTTAACAGCGATCGCGTATTTCGTCT<br>CGCTCAGGCGCAATCACGAATGAATAACGGTTTGGTTGATGCGAGTGATTTTGAT<br>GACGAGCGTAATGGCTGGCCTGTTGAACAAGTCTGGAAAGAAATGCATAAGCTTT<br>TGCCATTCTCACCGGATTCAGTCGTCACTCATGGTGATTTCTCACTTGATAACCT<br>TATTTTTGACGAGGGGAAATTAATAGGTTGTATTGATGTTGGACGTGTCGGAATC<br>GCAGACCGATACCAGGATCTTGCCATCCTATGGAACTGCCTCGGTGAGTTTTCTC<br>CTTCATTACAGAAACGGCTTTTTCAAAAATATGGTATTGATAATCCTGATATGAA<br>TAAATTGCAGTTTCATTTGATGCTCGATGAGTTTTTCTAAA | Young, 2018 | [1] |
| *chlR* | Cloning antibiotic cassette | gttctgaggtcattactggatctatcaacagcagtccaagcgagctcgatatcaa<br>attacgccccgccctgccactcatcgcagtactgttgtaattcattaagcattct<br>gccgacatggaagccatcacaaacggcatgatgaacctgaatcgccagcggcatc<br>agcaccttgtcgccttgcgtataatatttgcccatggtgaaaacgggggcgaaga<br>agttgtccatattggccacgtttaaatcaaaactggtgaaactcacccagggatt<br>ggctgagacgaaaaacatattctcaataaaccctttagggaaataggccaggttt<br>tcaccgtaacacgccacatcttgcgaatatatgtgtagaaactgccggaaatcgt<br>cgtggtattcactccagagcgatgaaaacgtttcagtttgctcatggaaaacggt<br>gtaacaagggtgaacactatcccatatcaccagctcaccgtctttcattgccata<br>cgaaattccggatgagcattcatcaggcgggcaagaatgtgaataaaggccggat<br>aaaacttgtgcttatttttctttacggtctttaaaaaggccgtaatatccagctg<br>aacggtctggttataggtacattgagcaactgactgaaatgcctcaaaatgttct<br>ttacgatgccattgggatatatcaacggtggtatatccagtgatttttttctcca<br>ttttagcttccTTAGCTCCTGAAAATCTCGATAACTCAAAAAATACGCCCGGTAG<br>TGATCTTATTTCATTATGGTGAAAGTTGGAACCTCTTACGTGCCCGATCAACTCG<br>CGCGTTTGCCACCTGACGTCTAAGAAAAGGAATATTCAGCAATTTGCCCGTGCCG<br>AAGAAAGGCCCACTG | Young, 2018 | [1] |
| *ampR* | Cloning antibiotic cassette | ttaccaatgcttaatcagtgaggcacctatctcagcgatctgtctatttcgttca<br>tccatagttgcctgactccccgtcgtgtagataactacgatacgggagggcttac<br>catctggccccagtgctgcaatgataccgcgagagccacgctcaccggctccaga<br>tttatcagcaataaaccagccagccggaagggccgagcgcagaagtggtcctgca<br>actttatccgcctccatccagtctattaattgttgccgggaagctagagtaagta<br>gttcgccagttaatagtttgcgcaacgttgttgccattgctacaggcatcgtggt<br>gtcacgctcgtcgtttggtatggcttcattcagctccggttcccaacgatcaagg<br>cgagttacatgatcccccatgttgtgcaaaaaagcggttagctccttcggtcctc<br>cgatcgttgtcagaagtaagttggccgcagtgttatcactcatggttatggcagc<br>actgcataattctcttactgtcatgccatccgtaagatgcttttctgtgactggt<br>gagtactcaaccaagtcattctgagaatagtgtatgcggcgaccgagttgctctt<br>gcccggcgtcaatacgggataataccgcgccacatagcagaactttaaaagtgct<br>catcattggaaaacgttcttcggggcgaaaactctcaaggatcttaccgctgttg<br>agatccagttcgatgtaacccactcgtgcacccaactgatcttcagcatctttta<br>ctttcaccagcgtttctgggtgagcaaaaacaggaaggcaaaatgccgcaaaaaa<br>gggaataagggcgacacggaaatgttgaatactcatactcttcctttttcaatat<br>tattgaagcatttatcagggttattgtctcatgagcggatacatatttgaatgta<br>tttagaaaaataaacaaataggggttccgcgcacatttccccgaaaagtgccacc<br>tgacgtctaagaaaccattattatcatgacattaacctataaaaataggcgtatc<br>acgaggccctttcgtctcgcgcgtttcggtgatgacggtgaaaacctctgacaca<br>tgcagctcccggagacggtcacagcttgtctgtaagcggatgccgggagcagaca<br>agcccgtcagggcgcgtcagcgggtgttggcgggtgtcggggctggcttaactat<br>gcggcatcagagcagattgtactg | Young, 2018 | [1] |

| | | | | |
|---|---|---|---|---|
| *vGFP* | Venus GFP gene | atgtctaaaggtgaagaattattcactggtgttgtcccaattttggttgaattag<br>atggtgatgttaatggtcacaaattttctgtctccggtgaaggtgaaggtgatgc<br>tacttacggtaaattgaccttaaaattgatttgtactactggtaaattgccagtt<br>ccatggccaaccttagtcactactttaggttatggtttgcaatgttttgctagat<br>acccagatcatatgaaacaacatgactttttcaagtctgccatgccagaaggtta<br>tgttcaagaaagaactatttttttcaaagatgacggtaactacaagaccagagct<br>gaagtcaagtttgaaggtgataccttagttaatagaatcgaattaaaaggtattg<br>attttaaagaagatggtaacattttaggtcacaaattggaatacaactataactc<br>tcacaatgtttacatcactgctgacaaacaaaagaatggtatcaaagctaacttc<br>aaaattagacacaacattgaagatggtggtgttcaattagctgaccattatcaac<br>aaaatactccaattggtgatggtccagtcttgttaccagacaaccattacttatc<br>ctatcaatctgccttatccaaagatccaaacgaaaagagagatcacatggtcttg<br>ttagaatttgttactgctgctggtattacccatggtatggatgaattgtacaaat<br>aa | Young, 2018 | [1] |

**Table S2:** Primers and oligos used in this study.

| Name | Type | Sequence | Source | Reference |
|---|---|---|---|---|
| crtI foward | qPCR primer | CAAGACCGACAGATTACGAAGAG | This study | |
| crtI rev | qPCR primer | CGGTGTATTGGAGCAAGGAATA | This study | |
| crtI probe | qPCR probe | /56-FAM/TACATGGGT/ZEN/CAAAGCCCATACAGTGC/3IABkFQ/ | This study | |
| crtS fwd | qPCR primer | CGTTAAGTCGATGGTCCCAAT | This study | |
| crtS rev | qPCR primer | CGAGTCTGGTTGCCTTCTTT | This study | |
| crtS probe | qPCR probe | /56-FAM/CAAGATGAT/ZEN/GGAGGATGCGGCTGA/3IABkFQ/ | This study | |
| FPS fwd | qPCR primer | CTCATCCCGATGGGTGAATAC | This study | |
| FPS rev | qPCR primer | ATGTCGGTTCCGATCTTTCC | This study | |
| FPS probe | qPCR probe | /56-FAM/AGTTCAGGA/ZEN/TGATGTGCTCGACGC/3IABkFQ/ | This study | |
| IDI fwd | qPCR primer | CGATCTGGTTCTCCTCTTCAAC | This study | |
| IDI rev | qPCR primer | GAAGAGCGGACGAGAAGATTAC | This study | |
| IDI probe | qPCR probe | /56-FAM/ACGTGTTGC/ZEN/AGTCATCCTTTGAGC/3IABkFQ/ | This study | |
| MVD fwd | qPCR primer | GATTTCGACTCGTTCGCAAAG | This study | |
| MVD rev | qPCR primer | CGGGAGACATCGTTCAAGTAA | This study | |
| MVD probe | qPCR probe | /56-FAM/TGAAGGACT/ZEN/CCAACCAGTTCCACG/3IABkFQ/ | This study | |
| PMVK fwd | qPCR primer | CTAGCAGGTGGATACCTTGTG | This study | |
| PMVK rev | qPCR primer | GAAGGCGGAAGAGATCGAATAA | This study | |

| | | | | |
|---|---|---|---|---|
| PMVK probe | qPCR probe | /56-FAM/CGTGGTTTC/ZEN/GACCTCATCCAGGTT/3IABkFQ/ | This study | |
| MVK fwd | qPCR primer | TCGGGATGAATCTGAGGTAGAG | This study | |
| MVK rev | qPCR primer | TTCGACTGTGTGTCGGTATTG | This study | |
| MVK probe | qPCR probe | /56-FAM/TCATCAAGC/ZEN/AAATCGTGAGGGTCCT/3IABkFQ/ | This study | |
| EY099 | Level 0 sequencing primer | CAGACAAGCCCGTCAGG | Young, 2018 | [1] |
| EY100 | Level 1 sequencing primer | GTTATCCCCTGATTCTGTGG | Young, 2018 | [1] |
| EY101 | Level 1 sequencing primer | ATTCAGCAATTTGCCCG | Young, 2018 | [1] |
| XD_crtYB_5'_F_1000 | Level 2 sequencing primer | CAGAAGATGGGTCCACCG | This study | |
| XD_crtYB_3'_R_1000 | Level 2 sequencing primer | CTCCATGCTGCTCTCGT | This study | |
| crtYB fwd | qPCR primer | CACCAAAGGAAGAGAGGATGAG | This study | |
| crtYB rev | qPCR primer | GTACTACCCATTGAGGAAGCTATG | This study | |
| crtYB probe | qPCR probe | /56-FAM/ATTGTTCTG/ZEN/GGTCTGTCTGCCTGG/3IABkFQ/ | This study | |
| crtE fwd | qPCR primer | GTGTCCTAGCGAAGAGGAATATG | This study | |
| crtE rev | qPCR primer | CCGAACATGGTCAGCAATAGA | This study | |
| crtE probe | qPCR probe | /56-FAM/TGGTTCTTG/ZEN/GAAGTGAGCGAATGGT/3IABkFQ/ | This study | |

**Table S3:** Strain modifications performed in this study. The parental strain CBS 6938 was first reported by Golubev, 1995 [5].

| Strain | Chromosome Insert | Plasmid Inserted |
|---|---|---|
| XdUV_1 | crtYB::($P_{HP1}$-*NLuc*-$T_{act}$//$P_{gdh}$-*natR*-$T_{gpd}$) | pJHCcrtYB1000:($P_{HP1}$-*NLuc*-$T_{act}$//$P_{gdh}$-*natR*-$T_{gpd}$//*kanR*//ColE1) |
| XdUV_2 | crtYB::($P_{Nth}$-*NLuc*-$T_{act}$//$P_{gdh}$-*natR*-$T_{gpd}$) | pJHCcrtYB1000:($P_{Nth}$-*NLuc*-$T_{act}$//$P_{gdh}$-*natR*-$T_{gpd}$ //*kanR*//ColE1) |
| XdUV_3 | crtYB::($P_{CCP}$-*NLuc*-$T_{act}$//$P_{gdh}$-*natR*-$T_{gpd}$) | pJHCcrtYB1000:($P_{CCP}$-*NLuc*-$T_{act}$//$P_{gdh}$-*natR*-$T_{gpd}$ //*kanR*//ColE1) |
| XdUV_4 | crtYB::($P_{TR}$-*NLuc*-$T_{act}$//$P_{gdh}$-*natR*-$T_{gpd}$) | pJHCcrtYB1000:($P_{TR}$-*NLuc*-$T_{act}$//$P_{gdh}$-*natR*-$T_{gpd}$ //*kanR*//ColE1) |
| XdUV_5 | crtYB::($P_{NUCR}$-*NLuc*-$T_{act}$//$P_{gdh}$-*natR*-$T_{gpd}$) | pJHCcrtYB1000:($P_{NUCR}$-*NLuc*-$T_{act}$//$P_{gdh}$-*natR*-$T_{gpd}$ //*kanR*//ColE1) |
| XdUV_6 | crtYB::($P_{CYC1}$-*NLuc*-$T_{act}$//$P_{gdh}$-*natR*-$T_{gpd}$) | pJHCcrtYB1000:($P_{CYC1}$-*NLuc*-$T_{act}$//$P_{gdh}$-*natR*-$T_{gpd}$ //*kanR*//ColE1) |
| XdUV_7 | crtYB::($P_{HAP3}$-*NLuc*-$T_{act}$//$P_{gdh}$-*natR*-$T_{gpd}$) | pJHCcrtYB1000:($P_{HAP3}$-*NLuc*-$T_{act}$//$P_{gdh}$-*natR*-$T_{gpd}$ //*kanR*//ColE1) |
| XdUV_8 | crtYB::($P_{GYG}$-*NLuc*-$T_{act}$//$P_{gdh}$-*natR*-$T_{gpd}$) | pJHCcrtYB1000:($P_{GYG}$-*NLuc*-$T_{act}$//$P_{gdh}$-*natR*-$T_{gpd}$ //*kanR*//ColE1) |
| XdUV_9 | crtYB::($P_{SAM}$-*NLuc*-$T_{act}$//$P_{gdh}$-*natR*-$T_{gpd}$) | pJHCcrtYB1000:($P_{SAM}$-*NLuc*-$T_{act}$//$P_{gdh}$-*natR*-$T_{gpd}$ //*kanR*//ColE1) |
| XdUV_10 | crtYB::($P_{HP2}$-*NLuc*-$T_{act}$//$P_{gdh}$-*natR*-$T_{gpd}$) | pJHCcrtYB1000:($P_{HP2}$-*NLuc*-$T_{act}$//$P_{gdh}$-*natR*-$T_{gpd}$ //*kanR*//ColE1) |
| XdUV_11 | crtYB::($P_{Con-6}$-*NLuc*-$T_{act}$//$P_{gdh}$-*natR*-$T_{gpd}$) | pJHCcrtYB1000:($P_{Con-6}$-*NLuc*-$T_{act}$//$P_{gdh}$-*natR*-$T_{gpd}$ //*kanR*//ColE1) |
| XdUV_12 | crtYB::($P_{SDR}$-*NLuc*-$T_{act}$//$P_{gdh}$-*natR*-$T_{gpd}$) | pJHCcrtYB1000:($P_{SDR}$-*NLuc*-$T_{act}$//$P_{gdh}$-*natR*-$T_{gpd}$ //*kanR*//ColE1) |
| XdUV_13 | crtYB::($P_{ETFQO}$-*NLuc*-$T_{act}$//$P_{gdh}$-*natR*-$T_{gpd}$) | pJHCcrtYB1000:($P_{ETFQO}$-*NLuc*-$T_{act}$//$P_{gdh}$-*natR*-$T_{gpd}$ //*kanR*//ColE1) |
| XdUV_14 | crtYB::($P_{MSS4}$-*NLuc*-$T_{act}$//$P_{gdh}$-*natR*-$T_{gpd}$) | pJHCcrtYB1000:($P_{MSS4}$-*NLuc*-$T_{act}$//$P_{gdh}$-*natR*-$T_{gpd}$ //*kanR*//ColE1) |
| XdUV_15 | crtYB::($P_{HP3}$-*NLuc*-$T_{act}$//$P_{gdh}$-*natR*-$T_{gpd}$) | pJHCcrtYB1000:($P_{HP3}$-*NLuc*-$T_{act}$//$P_{gdh}$-*natR*-$T_{gpd}$ //*kanR*//ColE1) |
| XdUV_16 | crtYB::($P_{E3}$-*NLuc*-$T_{act}$//$P_{gdh}$-*natR*-$T_{gpd}$) | pJHCcrtYB1000:($P_{E3}$-*NLuc*-$T_{act}$//$P_{gdh}$-*natR*-$T_{gpd}$ //*kanR*//ColE1) |
| XdUV_17 | crtYB::($P_{UspA}$-*NLuc*-$T_{act}$//$P_{gdh}$-*natR*-$T_{gpd}$) | pJHCcrtYB1000:($P_{UspA}$-*NLuc*-$T_{act}$//$P_{gdh}$-*natR*-$T_{gpd}$ //*kanR*//ColE1) |
| XdUV_18 | crtYB::($P_{Grg1}$-*NLuc*-$T_{act}$//$P_{gdh}$-*natR*-$T_{gpd}$) | pJHCcrtYB1000:($P_{Grg1}$-*NLuc*-$T_{act}$//$P_{gdh}$-*natR*-$T_{gpd}$ //*kanR*//ColE1) |

| | | |
|---|---|---|
| XdUV_19 | crtYB::($P_{ADH}$-*NLuc*-$T_{act}$//$P_{gdh}$-*natR*-$T_{gpd}$) | pJHCcrtYB1000:($P_{ADH}$-*NLuc*-$T_{act}$//$P_{gdh}$-*natR*-$T_{gpd}$ //*kanR*//ColE1) |
| XdUV_20 | crtYB::($P_{HP4}$-*NLuc*-$T_{act}$//$P_{gdh}$-*natR*-$T_{gpd}$) | pJHCcrtYB1000:($P_{HP4}$-*NLuc*-$T_{act}$//$P_{gdh}$-*natR*-$T_{gpd}$ //*kanR*//ColE1) |
| XdUV_21 | crtYB::($P_{RP}$-*NLuc*-$T_{act}$//$P_{gdh}$-*natR*-$T_{gpd}$) | pJHCcrtYB1000:($P_{RP}$-*NLuc*-$T_{act}$//$P_{gdh}$-*natR*-$T_{gpd}$ //*kanR*//ColE1) |
| XdUV_22 | crtYB::($P_{HP5}$-*NLuc*-$T_{act}$//$P_{gdh}$-*natR*-$T_{gpd}$) | pJHCcrtYB1000:($P_{HP5}$-*NLuc*-$T_{act}$//$P_{gdh}$-*natR*-$T_{gpd}$ //*kanR*//ColE1) |
| XdUV_23 | crtYB::($P_{HP6}$-*NLuc*-$T_{act}$//$P_{gdh}$-*natR*-$T_{gpd}$) | pJHCcrtYB1000:($P_{HP6}$-*NLuc*-$T_{act}$//$P_{gdh}$-*natR*-$T_{gpd}$ //*kanR*//ColE1) |
| XdUV_24 | crtYB::($P_{MLRQ}$-*NLuc*-$T_{act}$//$P_{gdh}$-*natR*-$T_{gpd}$) | pJHCcrtYB1000:($P_{MLRQ}$-*NLuc*-$T_{act}$//$P_{gdh}$-*natR*-$T_{gpd}$ //*kanR*//ColE1) |
| XdUV_25 | crtYB::($P_{RPL6}$-*NLuc*-$T_{act}$//$P_{gdh}$-*natR*-$T_{gpd}$) | pJHCcrtYB1000:($P_{RPL6}$-*NLuc*-$T_{act}$//$P_{gdh}$-*natR*-$T_{gpd}$ //*kanR*//ColE1) |
| XdUV_26 | crtYB::($P_{RPL19}$-*NLuc*-$T_{act}$//$P_{gdh}$-*natR*-$T_{gpd}$) | pJHCcrtYB1000:($P_{RPL19}$-*NLuc*-$T_{act}$//$P_{gdh}$-*natR*-$T_{gpd}$ //*kanR*//ColE1) |

**Table S4:** Name determination through consensus.

| Scaffold Location | ERGO Gene Ontology | Interpro Annotation | JGI Prediction | BLAST | Consensus Name | Promoter Name (this work) |
|---|---|---|---|---|---|---|
| scaffold_3: 894515..896377 | Molecular Function Biological Process response to stress<br><br>Cellular Component | IPR006016 UspA | Hypothetical protein | Universal stress protein A (UspA) | Universal stress protein A (UspA) | $P_{UspA}$ |
| scaffold_2: 1922200 ..1922852 | None provided | IPR020100 Glucose-repressible protein Grg1 | Glucose repressible protein 2 | Glucose repressible protein 2 | Glucose repressible protein 2 | $P_{Grg1}$ |
| scaffold_10: 164277..165920 | Molecular Function alcohol dehydrogenase (NAD+) activity<br><br>alcohol dehydrogenase activity, metal ion-independent<br><br>alcohol dehydrogenase activity, zinc-dependent<br><br>alcohol dehydrogenase activity, iron-dependent<br><br>Biological Process Cellular Component | (unassigned)(EC 1.1.1.1)<br><br>IPR011032 GroES (chaperonin 10)-like<br><br>IPR020843 Polyketide synthase, enoylreductase domain<br><br>IPR013154 Alcohol dehydrogenase GroES-like<br><br>IPR013149 Alcohol dehydrogenase, C-terminal | Alcohol dehydrogenase | Alcohol dehydrogenase | Alcohol dehydrogenase | $P_{ADH}$ |
| scaffold_1: 1665495..1666335 | None provided | IPR022234 Protein of unknown function DUF3759 | Protein of unknown function | Phosphoglycerate mutase family | Hypothetical protein 4 | $P_{HP4}$ |
| scaffold_9: 680348..682293 | None provided | IPR007946 A1 cistron-splicing factor, AAR2 | Putative ubiquitin/ribosomal protein S27a fusion protein | Ubiquitin-40s ribosomal protein s31 fusion protein | Ribosomal protein (RP) | $P_{RP}$ |

| | | | | | | |
|---|---|---|---|---|---|---|
| scaffold_14: 331267..331837 | None provided | None provided | Expressed protein | Hypothetical protein | Hypothetical protein 5 | $P_{HP5}$ |
| scaffold_4: 316497..318032 | None provided | None provided | Hypothetical protein | Hypothetical protein | Hypothetical protein 6 | $P_{HP6}$ |
| scaffold_5: 1150021..1150603 | None provided | IPR010530 NADH-ubiquinone reductase complex 1 MLRQ subunit | Hypothetical protein | NADH-ubiquinone reductase complex 1 MLRQ subunit | NADH-ubiquinone reductase complex 1 MLRQ subunit | $P_{MLRQ}$ |
| scaffold_2: 1298264..1299037 | Molecular Function structural constituent of ribosome<br><br>Biological Process translation<br><br>Cellular Component intracellular anatomical structure<br><br>ribosome | IPR000702 Ribosomal protein L6<br><br>IPR020040 Ribosomal protein L6, alpha-beta domain | Ribosomal protein L6%2C alpha-beta domain-containing protein | Glucosyltransferase-Alg8p | Ribosomal protein L6, alpha-beta domain | $P_{RPL6}$ |
| scaffold_8: 84498..85356 | Molecular Function structural constituent of ribosome<br><br>Biological Process translation<br><br>Cellular Component intracellular anatomical structure<br><br>ribosome | IPR023638 Ribosomal protein L19/L19e conserved site<br><br>IPR000196 Ribosomal protein L19/L19e domain | 60S ribosomal protein L19 | 60S ribosomal protein L19 | 60S ribosomal protein L19 | $P_{RPL19}$ |
| scaffold_1: 1544039..1548049 | Molecular Function protein binding<br><br>zinc ion binding<br><br>ATP-dependent peptidase activity | IPR015947 PUA-like domain<br><br>IPR001841 Zinc finger, RING-type<br><br>IPR017907 Zinc finger, RING-type, conserved site | Hypothetical protein | Predicted E3 ubiquitin ligase | Predicted E3 ubiquitin ligase | $P_{E3}$ |

| | | | | | | |
|---|---|---|---|---|---|---|
| | Biological Process obsolete ATP-dependent proteolysis | IPR003111 Peptidase S16, lon N-terminal | | | | |
| scaffold_2: 2209212..2210525 | None provided | None provided | Hypothetical protein | Hypothetical protein | Hypothetical protein 3 | $P_{HP3}$ |
| scaffold_2: 1904960..1906561 | None provided | None provided | Hypothetical protein | Hypothetical protein | Hypothetical protein 2 | $P_{HP2}$ |
| scaffold_3: 348899..350906 | None provided | None provided | Expressed protein | Hypothetical protein | Hypothetical protein 1 | $P_{HP1}$ |
| scaffold_5: 1538428..1540736 | Molecular Function catalytic activity<br><br>DNA-(apurinic or apyrimidinic site) endonuclease activity<br><br>Biological Process base-excision repair<br><br>DNA repair | (unassigned) (EC 4.2.99.18)<br><br>IPR011257 DNA glycosylase<br><br>IPR003265 HhH-GPD domain | DNA glycosylase | NTH | Endonuclease III | $P_{Nth}$ |
| scaffold_7: 1106783..1108757 | Molecular Function hexosyltransferase activity<br><br>inositol 3-alpha-galactosyltransferase activity<br><br>Biological Process carbohydrate biosynthetic process | (unassigned) (EC 2.4.1.123)<br><br>IPR029044 Nucleotide-diphospho-sugar transferases<br><br>IPR002495 Glycosyl transferase, family 8 | Nucleotide-diphospho-sugar transferase | Glycosyltransferase family 8 protein | Glycogenin glucosyltransferase | $P_{GYG}$ |
| scaffold_8: 1138346..1140885 | Molecular Function thiopurine S-methyltransferase activity<br><br>Biological Process metabolic process | IPR029063 S-adenosyl-L-methionine-dependent methyltransferase<br><br>IPR008854 TPMT family | S-adenosyl-L-methionine-dependent methyltransferase | TPMT family | S-adenosyl-L-methionine-dependent methyltransferase | $P_{SAM}$ |

| | | | | | | |
|---|---|---|---|---|---|---|
| | Cellular Component cytoplasm | | | | | |
| scaffold_5: 138744..140405 | Molecular Function carbon-sulfur lyase activity<br><br>Biological Process metabolic process | IPR011057 Mss4-like<br><br>IPR006913 Glutathione-dependent formaldehyde-activating enzyme/centromere protein V | Mss4-like protein | Mss4-like | MSS4-like protein | $P_{MSS4}$ |
| scaffold_12: 632916..634567 | Molecular Function oxidoreductase activity<br><br>glucose dehydrogenase activity<br><br>pinocarveol dehydrogenase activity<br><br>chloral hydrate dehydrogenase activity<br><br>hydroxymethylmethylsilanediol oxidase activity<br><br>1-phenylethanol dehydrogenase activity<br><br>myrtenol dehydrogenase activity<br><br>versicolorin reductase activity<br><br>ketoreductase activity<br><br>Biological Process metabolic process | (unassigned) (EC 1.1.-.-)<br><br>IPR002198 Short-chain dehydrogenase/reductase SDR | putative short-chain dehydrogenase TIC 32%2C chloroplastic | Short-chain dehydrogenase | Short-chain dehydrogenase | $P_{SDR}$ |
| scaffold_2 :1824823..1825250 | None provided | IPR018824 Conidiation-specific protein 6 | Hypothetical protein | Conidiation-specific protein 6 | Conidiation-specific protein 6 | $P_{Con-6}$ |

| | | | | | | |
|---|---|---|---|---|---|---|
| scaffold_11: 37744..38752 | Molecular Function iron ion binding<br><br>electron transfer activity<br><br>heme binding<br><br>Biological Process obsolete electron transport | IPR009056 Cytochrome c-like domain | cytochrome c C1 | Cytochrome c1 | Cytochrome c1 | $P_{CYC1}$ |
| scaffold_8: 140491..141329 | Molecular Function sequence-specific DNA binding<br><br>DNA binding<br><br>DNA-directed DNA polymerase activity<br><br>DNA polymerase activity<br><br>Biological Process Cellular Component intracellular anatomical structure | (unassigned) (EC 2.7.7.7)<br><br>IPR009072 Histone-fold<br><br>IPR003958 Transcription factor CBF/NF-Y/archaeal histone | AGAP008344-PA | Transcription factor hap3 | Transcription activator HAP3 | $P_{HAP3}$ |
| scaffold_1: 1018792..1022011 | Molecular Function S-adenosylmethionine-dependent tRNA (m5U54) methyltransferase activity<br><br>Biological Process Cellular Component | (unassigned) (EC 2.1.1.35)<br><br>IPR029063 S-adenosyl-L-methionine-dependent methyltransferase<br><br>IPR025795 tRNA (uracil-5-)-methyltransferase<br><br>IPR030391 RNA methyltransferase TrmA, conserved site<br><br>IPR030390 RNA methyltransferas | putative tRNA methyltransferase | tRNA methyltransferase | Endo-exonuclease NUCR | $P_{NUCR}$ |

| | | | | | | |
|---|---|---|---|---|---|---|
| | | e TrmA, active site<br><br>IPR010280 (Uracil-5)-methyltransferase family | | | | |
| scaffold_4: 725147..726268 | Molecular Function FMN binding | IPR012349 FMN-binding split barrel<br><br>IPR007396 Transcriptional regulator PAI 2-type | Putative FMN-binding domain-domain containing protein | Transcriptional regulator | Transcriptional regulator | $P_{TR}$ |
| scaffold_1: 2045089..2048287 | Molecular Function electron-transferring-flavoprotein dehydrogenase activity<br><br>Biological Process obsolete electron transport | (unassigned) (EC 1.5.5.1)<br><br>IPR017896 4Fe-4S ferredoxin-type, iron-sulphur binding domain<br><br>IPR007859 Electron transfer flavoprotein-ubiquinone oxidoreductase | putative oxidoreductase | Electron-transferring-flavoprotein dehydrogenase | Electron transfer flavoprotein-ubiquinone oxidoreductase | $P_{ETFQO}$ |
| scaffold_6: 802427..804411 | Molecular Function peroxidase activity<br><br>heme binding<br><br>cytochrome-c peroxidase activity<br><br>Biological Process obsolete electron transport<br><br>response to oxidative stress | (unassigned) (EC 1.11.1.5)<br><br>IPR010255 Haem peroxidase<br><br>IPR019794 Peroxidase, active site<br><br>IPR019793 Peroxidases heam-ligand binding site<br><br>IPR002016 Haem peroxidase, plant/fungal/bacterial | mitochondrial Cytochrome c peroxidase | Cytochrome c peroxidase | Cytochrome c peroxidase | $P_{CCP}$ |

### SUPPLEMENTARY FIGURES

a.

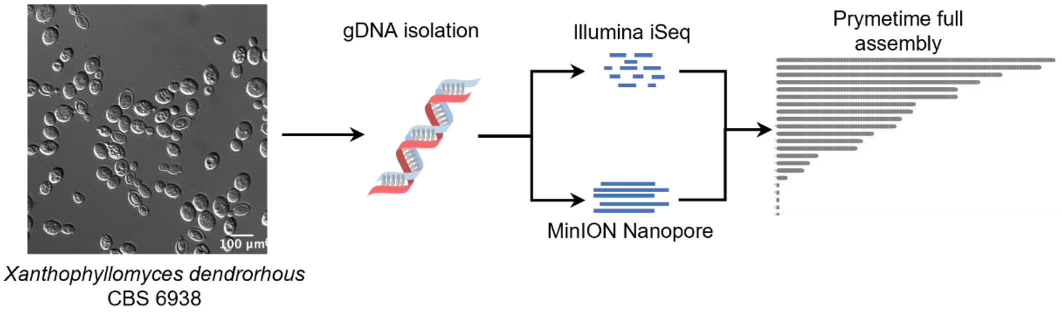

b.

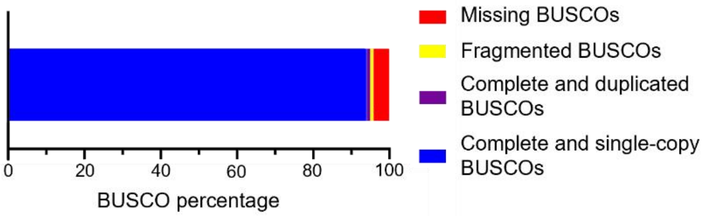

**Figure S1:** ***X. dendrorhous*** **CBS 6398 genome sequencing. (a)** Schematic of Prymetime next generation sequencing workflow. The left is a brightfield microscope image of *X. dendrorhous*, the middle describes genomic DNA isolation and they hybrid sequencing approach of Prymetime. The image on the right is a Chromomap depiction of the completed resequenced *X. dendrorhous* genome with 16 large contigs and several smaller fragments. **(b)** BUSCO analysis of the *X. dendrorhous* genome depicting the 94% completeness.

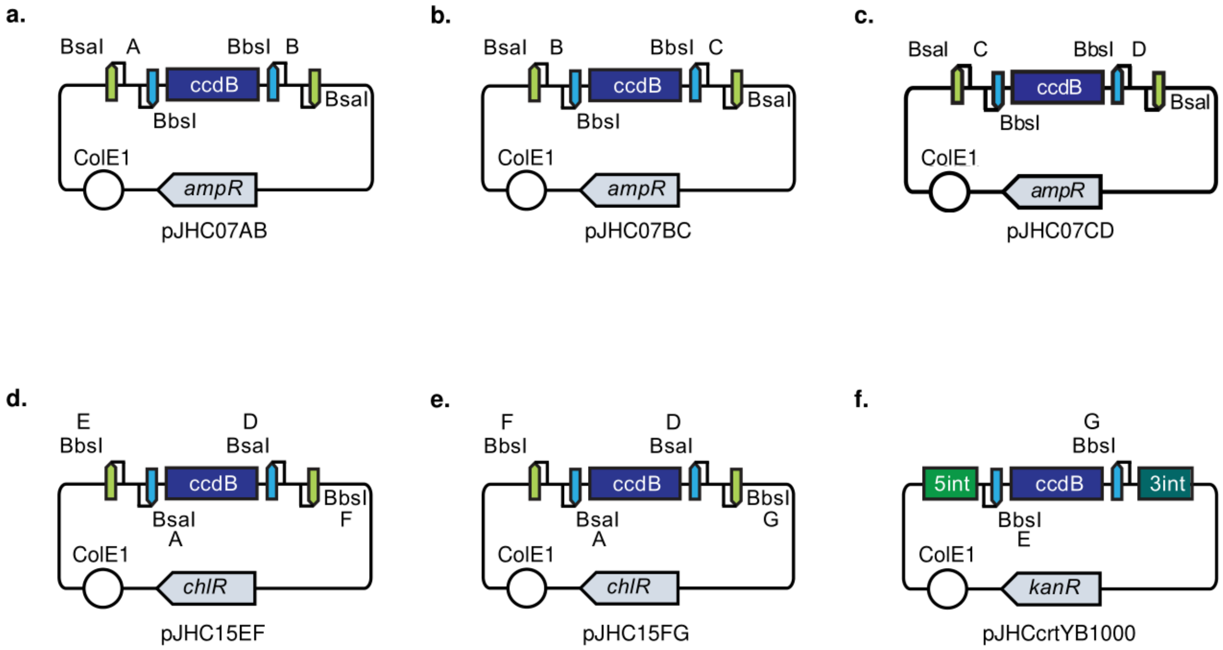

**Figure S2: Plasmid maps used for cloning in this study.** All cloning vectors use ccdB knockout alongside antibiotic resistance as selection markers. **(a)** pJHC07AB is an L0 cloning vector used to propagate promoter sequences with A (GTGC) and B (AATG) scars. Both BbsI and BsaI sites have A and B scars. The plasmid contains the *ampR* gene which confers ampicillin resistance. **(b)** pJHC07BC is identical to pJHC07AB except for the scars B and C (TAAA). pJHC07BC is used to propagate ORF sequences. **(c)** pJHC07CD is identical to pJHC07AB except for the scars C and D (CCTC). pJHC07CD is used to propagate terminator sequences. **(d)** The L1 plasmid pJHC15EF is used to propagate transcription units combined from pJHC07AB, pJHC07BC, and pJHC07CD (promoter-ORF-terminator). After BsaI digestion and Type IIS reaction, the transcription unit is nested in BbsI restriction enzyme sites with E (GCTT) and F (CTGA) scars. The plasmid contains the *chlR* gene which confers chloramphenicol resistance. **(e)** The L1 plasmid pJHC15FG is identical to pJHC15EF except for the scars F and G (TACG). **(f)** pJHCcrtYB1000 is an L2 plasmid used to propagate transcription units from pJHC15EF and pJHC15FG that are joined by a Type IIS reaction. The BbsI sites are flanked by 1000 bp homology arms for integration into the crtYB site in *X. dendrorhous*. The plasmid contains the *kanR* gene which confers kanamycin resistance.
